# T-type calcium channels participate in intrinsic and synaptic activity of PKCγ neurons of the dorsal horn of the spinal cord during chronic pain

**DOI:** 10.1101/2024.12.11.627933

**Authors:** Célia Cuculière, Rachel Bourdon-Alonzeau, Coline Rulhe, Jean Chemin, Amaury François, Marie-Pierre Blanchard, Pierre Fontanaud, Emmanuel Deval, Matteo Mangoni, Eric Lingueglia, Emmanuel Bourinet, Pierre-François Méry

## Abstract

The disinhibition of the excitatory PKCγ interneurons plays a central role during mechanical allodynia in the dorsal horn of the spinal cord, routing harmless information to nociceptive pathways. The T-type calcium channel Cav3.2, necessary for mechanical and cold allodynia, is found in most PKCγ neurons of the spinal cord. In this study, the role of Cav3.2 in PKCγ neurons was studied after its pharmacological inhibition and its conditional deletion (KO) in Cav3.2^GFP-Flox^ KI x PKCγ-CreERT2 x Ai14 mice in normal conditions and in the spared-nerve-injury (SNI) model of neuropathic pain. Conditional deletion of Cav3.2 increased the hind-paw basal mechanical sensitivity before surgery, and decreased mechanical pain 7 days, but not 28 days, after surgery. At the cellular level, Cav3.2 participated in the low-threshold currents of PKCγ neurons and the T-type calcium current of PKCγ neurons was decreased in KO mice as compared to wild-type (WT). This loss did not convert into proportional alterations in subthreshold properties including “rebound” potentials, suggesting the involvement of other T-type channels. Action potential kinetics and firing properties seemed similar in WT and KO mice too, but rebound potentials were diminished in the SNI model in WT but not in KO mice. In addition, the modulations of firing properties induced by T-type channel pharmacological blocker Z944 observed in WT mice were absent in KO mice and after SNI. Furthermore, the pairing of action potentials was modified after SNI in WT mice, and not in KO mice. At the synaptic level, excitatory currents were lowered 7- and 28-days after surgery, while inhibitory currents were lowered only at 28 days. These changes were not found in Cav3.2-ablated neurons. Miniature currents analysis indicated that Cav3.2 was involved in both excitatory and inhibitory synaptic transmissions at the level of PKCγ neurons. Surprisingly, Z944 did not mimic the effects of Cav3.2 ablation in PKCγ neurons, suggesting distinct and eventually opposite roles of other T-type calcium channels. Altogether, our results show that Cav3.2 is not mandatory for firing of PKCγ neurons of the dorsal horn of the spinal cord, but that it participates to the SNI-induced changes in their intrinsic and synaptic activity, including changes in their excitatory and inhibitory controls.

**Graphical abstract:** 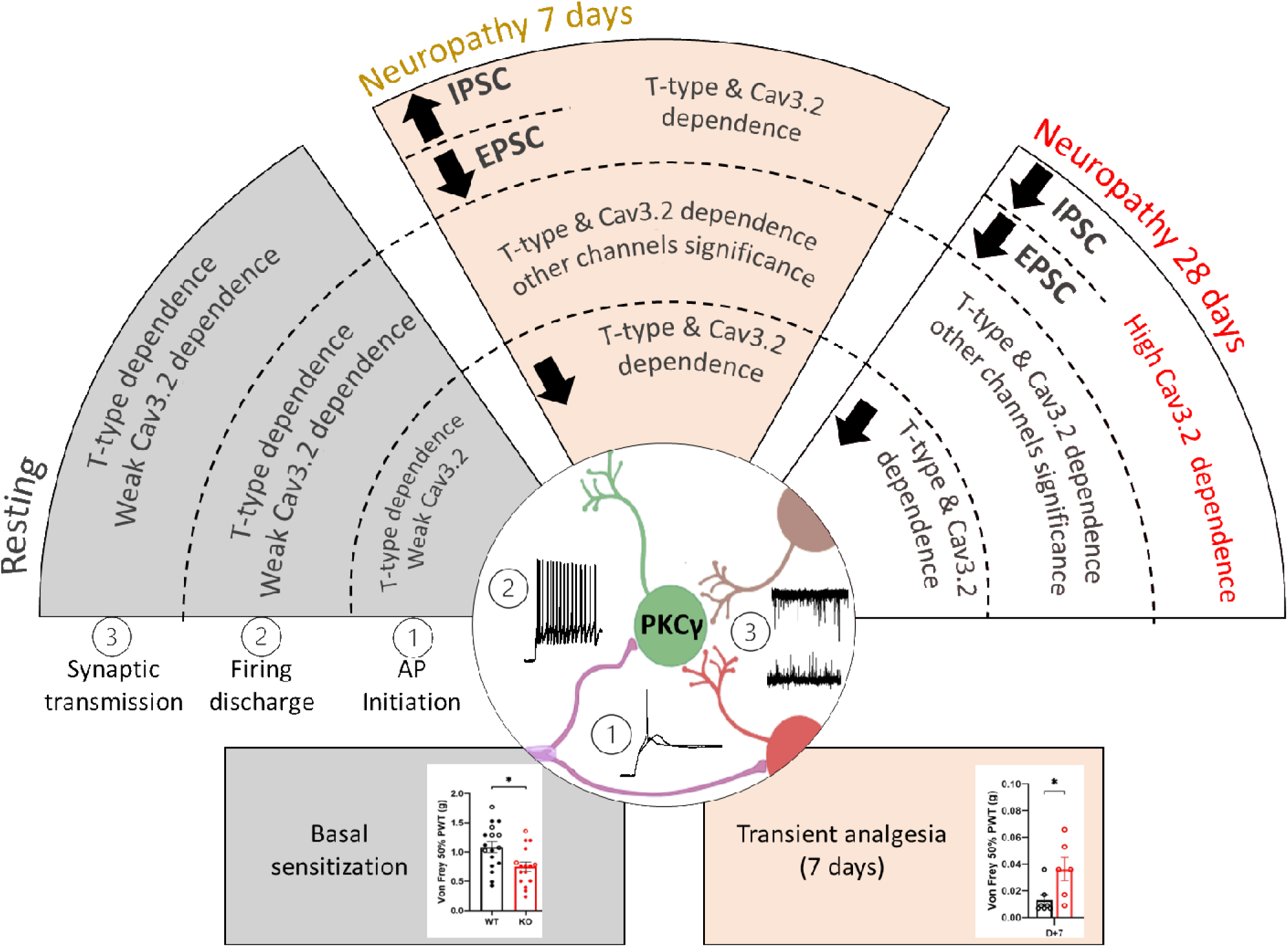

## Introduction

Chronic pain arises and progresses as a result of cellular remodelings within brain networks (Bardoni et al., 2013; Ghazisaeidi et al., 2023; Sandkühler, 2009). The spinal cord is well-known in undergoing major adaptations during chronic pain, broadly classified as disinhibitions and sensitizations. According to the gate control theory of pain, these remodeling are taking place within neuronal ensembles in the dorsal horn of the spinal cord. Indeed, during mechanical allodynia, the inhibitory parvalbumin neurons no longer dampen the activation of excitatory PKCγ neurons upon non noxious stimulation by low threshold primary afferent neurons (Lu et al., 2013; Neumann et al., 2008; Peirs et al., 2021; Petitjean et al., 2015; Qiu et al., 2022; Q. Wang et al., 2020). As a result, sensory impulses unrelated to nociceptive primary afferent neurons, are delivered to ascending nociceptive pathways (Bardoni et al., 2013; Peirs et al., 2021). Restoring an inhibitory control, selectively in PKCγ neurons, is sufficient in blunting mechanical allodynia and normalizing sensory transduction, in a mouse model of chronic pain (Peirs et al., 2015). This is supporting a strategy whereby the selective targeting of endogenous proteins expressed by PKCγ neurons might restore the inhibition of PKCγ neurons during allodynia and alleviate chronic pain. The low-threshold T-type, calcium channel Cav3.2, is homogenously expressed at the mRNA level (Russ et al., 2021) and is homogenously observed at the protein level (Candelas et al., 2019) in PKCγ neurons in mouse. NucSeq analysis shows that Cav3.2 is expressed in most PKCγ neurons in humans as well (Yadav et al., 2023). The intrathecal injection of cacna1h antisense shRNA blunts visceral pain (Marger et al., 2011), has an anti-allodynic effect in a neuropathic pain model in rats (Bourinet et al., 2005) and an anti-hyperalgesic effect in diabetic rats (Messinger et al., 2009). In addition, Cre-recombinase dependent RNA interference targeting of cacna1h in somatostatin-expressing interneurons in the spinal cord also has anti allodynic effects in mice (Zhi et al., 2022). Importantly, Cav3.1 and Cav3.3 do not share the same analgesic properties (Bourinet et al., 2005, 2016). This might be surprising since many neurons in the spinal cord, including PKCγ neurons, express more than one T-type calcium channel isoform (Candelas et al., 2019; Russ et al., 2021), which have few dissimilar biophysical properties (Zamponi et al., 2015). In other brain structures where differential roles for T-type channels were found, these were supported by differential targeting to selective subcellular compartments supporting different functions (Allken et al., 2014; Isope et al., 2012; Kovács et al., 2010; Lipkin et al., 2021; Molineux et al., 2006). Therefore, it seems important to decipher the role of Cav3.2 in PKCγ neurons.

While T-type calcium channels antagonists induce various forms of analgesia (Harding et al., 2021; Kerckhove et al., 2014; Nam, 2018; Shen et al., 2015; Snutch & Zamponi, 2018), none of them proved reliable in alleviating chronic pain, including neuropathic pain in humans. The pharmacology of T-type channels remains perfectible, and cell-type specific genetic manipulation in mouse models of chronic pain is an invaluable approach in evaluating the role of Cav3.2 in specific neurons, including PKCγ neurons.

The roles of Cav3.2 were resolved using Cav3.2^GFP-Flox^ KI mice in primary afferent neurons (François et al., 2015), in interneurons of the dorsal horn of the spinal cord (Candelas et al., 2019), and in parvalbumin-expressing GABAergic neurons of the thalamus (Fayad et al., 2022). Here, we examined PKCγ neurons in PKCγ-Cre^ERT2^ x Ai14 line (Abraira et al., 2017; Madisen et al., 2010). The selective ablation of cacna1h in PKCγ neurons was achieved by activating the tamoxifen-dependent Cre recombinase of the PKCγ-CreERT2 line with a homozygous Cav3.2^GFP-Flox^ KI background. We found that the selective deletion of Cav3.2 channels in PKCγ neurons has an antalgic effect at an early, but not late, time point of a chronic pain condition, elicited in the mouse by the spared nerve injury (Decosterd & Woolf, 2000). At the cellular level, deletion of Cav3.2 channels in PKCγ neurons counteracted some changes in the firing patterns induced by SNI in PKCγ neurons. In addition, deletion of Cav3.2 channels in PKCγ neurons counteracted many of the SNI-induced changes of the excitatory and the inhibitory controls of PKCγ neurons. Collectively, our findings reveal critical functions for Cav3.2 in the intrinsic activity as well as the synaptic activity of PKCγ neurons.

## MATERIALS AND METHODS

### Study approval

Animal procedures (#2016012216345875, #2021012115593979) complied with the welfare guidelines of the European Community and were ethically approved by the Direction of Veterinary departments of Herault, France (agreement number A 34-172-41).

### Animal models

The Cav3.2^GFP-Flox^ KI (François et al., 2015), PKCγ-Cre^ERT2^ (Abraira et al., 2017), and Ai14 (Madisen et al., 2010).knockin mouse lines were studied alone and in combination as described. Knocking out the Cav3.2 gene was achieved by activating the tamoxifen-dependent Cre recombinase of the PKCγ-CreERT2 line under a homozygous Cav3.2^GFP-Flox^ KI background. The treatment with tamoxifen (dissolved in ethanol and mixed with 9 volumes of kolliphor EL oil) was achieved by intraperitoneal injection (100µl at 10mg/ml) under isoflurane anaesthesia during 5 consecutive days, after the 4^th^ post-natal week.

*SNI surgery* method was adapted from (Decosterd & Woolf, 2000). Mice were anesthetized with isoflurane (4% induction; 2% steady-state) and were given meloxicam (Metacam 60µl, 1mg/ml i.p., 10min before skin incision). The skin on the lateral surface of the thigh was incised after multiple local injections of lidocaine (Laocaïne, 32µg/ml). The biceps femoris muscle exposing and separated allowing access to sciatic nerve and its three terminal branches: the sural, common peroneal and tibial nerves. The common peroneal and the tibial nerves were tight-ligated with 6-0 silk (Ethicon) and sectioned distal to the ligation, removing 2±4 mm of the distal nerve stump. Care was taken to avoid contact with or stretching of the intact sural nerve. After surgery, muscles were replaced and the skin was sutured before the mouse was transferred into a recovery cage for >4hours, and postsurgical treatments were provided according to guidelines.

### Behavioral tests

They were performed on 5 to 16 weeks-old mice in the SPF animal facility at room temperature (22-24°C) during the diurnal period of the circadian cycle (7:00-18:00). Mice were acclimatised for 20 min to their environment prior testing (François et al., 2015) and pain scores were determined in compliance with ethical guidelines (Jaggi et al., 2011). Static mechanical allodynia was measured using the Von Frey test. The animals were placed in Plexiglas chambers (8 x 8 cm) on an elevated grid and the arch of the posterior paw was tested using the up-down method (Chaplan et al., 1994).

### Slice preparation for electrophysiological recordings

Adult 6-12 weeks-old Ai14-positive mice of the appropriate genotype were anesthetized by injection of 0.1ml/10mg ketamine (10mg/ml) plus xylazine (1mg/ml). They were perfused through the left ventricle with solution-1 [in mM; 122 N-methyl-D-glucamine-Cl, 2.5 KCl, 1 CaCl_2_, 7 MgCl_2_, 2.5 NaHCO_3_, 1.5 NaH_2_PO_4_, 10 HEPES, 25 glucose, 3 Na-pyruvate, 2.5 kynurenic acid, 5 ascorbic acid, 0.2-5 thiourea, 2 N-acetyl-L-cysteine; pH 7.4, 0-2°C, gassed with O_2_] (Baccam et al., 2007). The lumbar region of the spinal cord was quickly removed in solution-1, embedded into low-melting Agarose (Invitrogen, Carlsbad, CA) and 300µm-thick frontal sections were processed on a microtome (Leica VT12000S, Leica Microsystem, Nanterre, France) within 20 min and kept at least for 1 hour in solution-2 [in mM; 115 NaCl, 2.5 KCl, 2 CaCl_2_, 4 MgCl_2_, 26 NaHCO_3_, 1.25 NaH_2_PO_4_, 25 glucose, 0.2 thiourea; pH 7.4, 30 °C, gassed with 95% CO_2_-5% O_2_].

### Patch-clamp recordings

Slices were immobilized with a nylon grid in a submersion chamber on the stage of an upright microscope (Olympus BX51WIF, Olympus Fr., Rungis, France) and superfused with solution-3 [in mM; 125 NaCl, 2.5 KCl, 2 CaCl_2_, 1 MgCl_2_, 26 NaHCO_3_, 1.25 NaH_2_PO_4_, 11 glucose; pH 7.4, 32°C, gassed with 95% CO_2_-5% O_2_] at 2 ml/min for at least 15 min. Infrared differential interference contrast illumination was used to visualize neurons, with a x40 immersion objective and Nomarski differential interference contrast optics, and the images captured with a camera (Jenoptik ProgRes MF, Bayeux, France). When appropriate, the tdTomato fluorescent signal was superimposed with the IR-DIC image in order to locate the PKCγ neurons. Borosilicate glass pipettes (6–8 MΩ) were connected to the head stage of an Axon MultiClamp 700B, interfaced with an Axon Digidata 1550A and controlled with pClamp10 (all from Molecular Devices, Sunnyvale, USA). Data were sampled at 20kHz and filtered at 3 kHz. Drugs were either bath-applied, by switching the supply of the superfusion system. Slices were discarded after being exposed to a compound. All chemicals were from Sigma-Aldrich (L’isle d’Abeau, France) excepted Z944 kindly provided by Dr. Terry Snutch (Michael Smith Laboratories, University of Columbia, Canada).

Whole cell patch-clamp experiments were performed with an internal medium (in mM): 139 K-gluconate, 10 HEPES, 0.1 EGTA acid, 1 MgCl_2_, 2 MgATP, 0.5 Na-GTP, 5 Na_2_-phospocreatine, 2.5 Na-pyruvate, 2 malate, pH 7.3 with KOH (295 mOsm adjusted with KMeSO_3_). Action potentials and resting properties were elicited by increasing or decreasing 2s-duration current pulses from a holding potential of -70mV (Grudt & Perl, 2002). Subthreshold electrical activity was triggered after a conditioning hyperpolarization at -90mV for 3s by step-by-step current increments of 1.5s-duration (Tadros et al., 2012). Transient sags during the conditioning step were taken into account when >3mV. Spontaneous excitatory and inhibitory currents were recorded at a steady-state potential of -70mV and -30mV, respectively. Miniature currents were recorded in the presence of extracellular tetrodotoxin (TTX) 1µM (Latoxan). Analyzes were performed with Axograph (Clements & Bekkers, 1997) or Clampex10 (Molecular Devices) using either a threshold based detection or an appropriate template. Action potential parameters were identified after peak detection with a Matlab standalone executable application (The Matworks Inc., Natick, MA). Firing patterns were classified as described (Candelas et al., 2019): Single spiking: less than 2 action potentials per pulse; Transient: activity duration < 1.4s in a 2s-long pulse; Irregular Tonic: tonic and having a SEM of the first 4 action potential intervals >2; Regular Tonic: tonic and having a SEM of the first 4 action potential intervals <2; Delayed: the latency of the first action potential >95ms and a mean frequency of the first 5 actions potentials >8Hz and Gap: the latency of the first action potential >95ms and a mean frequency of the first 5 action potentials <8Hz.

*For T-type calcium current recordings,* media compositions were adapted from Dreyfus et al. (2010). The internal solution was (in mM): 125 CsMeSO_3_, 10 HEPES, 5 EGTA acid, 1 MgCl_2_, 2 MgATP, 0.5 Na-GTP, 5 Na_2_-phospocreatine, 2 Na-pyruvate, 2 malate, 7 KCl, pH 7.4 with CsOH (295 mOsm adjusted with CsMeSO_3_). The external medium contained (in mM): 85 NaCl, 10 D-glucose, 10 CsCl, 2.5 KCl, 2 CaCl_2_, 1 MgCl_2_, 2,5 NaHCO_3_, 1.25 NaH_2_PO_4_, 10 HEPES, 5 4-aminopyridine (4-AP), 40 TEA-Cl, pH 7.4, 32°C, gassed with O_2_. After breaking the patch, blockers (TTX 1µM, APV 50µM, DNQX 30µM, GABAzine 3µM, strychnine 10µM) were added to the superfusion medium, and T-type channel blockers (NiCl_2_, ZnCl_2_, Z944) were further added at steady-state. A routine 2-steps protocol (-45mV; 250ms / -30mV; 250 ms) was elicited every 10s from a holding of -90mV. For current-voltage relationships, neurons were held at -80 mV and every 8 seconds, a 750ms-long conditioning pulse at -100mV preceded the 250-ms long test-pulse at 80mV, which was increased by steps of 5mV, ended by repolarization to -50mV for 250ms. T type calcium current was calculated as the difference between the maximal peak amplitude and the end pulse current.

### Immunofluorescence experiments

Mice (8-14 weeks) were processed according to Candelas et al. (2019) after anesthesia with ketamine plus xylazine. Tissues were fixed with a transcardiac perfusion of cold PBS containing PFA 4% and a post-fixation (2 hours) at 4°C. Samples were either embedded in 4% agarose and sectioned at 60 µm with a vibratome (Microm HM 650V, Brignais, France) or included in Tissue Tek OCT compound (Sakura Finetek, USA) and sectioned at 16µm with a cryostat (Microm NX70DV). The sections were blocked with TBS plus 0.05% tween 20 (TBST), 0.2% Triton X 100 and 10% normal serum (goat or donkey), 22°C, for 1 hr; incubations with primary antibodies (**Table 1**) were done in blocking solution at 4°C overnight; followed after 3x washes in PBS by an incubation with secondary antibodies (**Table 1**) were performed at RT for 2 hrs. They were included in Dako Mounting Medium (Agilent Tech., Santa Clara, USA) for imaging with a confocal microscope (Leica SP8-UV, Nanterre, France) equipped with 20X (Plan Apochromat 0.75 NA Imm Corr) or x40 (Plan Apochromat 1.3 NA oil DIC) objectives. Image processing was performed with FiJi (Schindelin et al., 2012), and cell densities were normalized to DAPI labelling or to the appropriate antigen using its Cell Counter plugin (Kurt DeVos, University of Sheffield, Academic Neurology).

**Table 1.** Antibodies & Probes.

| **Antibodies and probes** |  |  |  |
| --- | --- | --- | --- |
| **Antigen** | **Species** | **Dilution** | **Provider** |
| **GFP** | Rabbit | 1/500 | Chromotek |
| **PKCγ** | Guinea pig | 1/800 | Frontier Institute |
| **Ankyrin G** | Mouse | 1/350 | Neuromab |
| **α-rabbit** | Goat | 1/500 | Invitrogen |
| **α-guinea pig** | Goat | 1/500 | Invitrogen |
| **α-mouse** | Goat | 1/500 | Invitrogen |
| **α-rabbit** | Goat | 1/500 | Jackson |
| **α-guinea pig** | Goat | 1/500 | Invitrogen |
| **Target** | **Species** | **Reference** | **Provider** |
| **CACNA1H** | Mouse | 459751-C3 | Biotechne |
| **CACNA1G** | Mouse | 459761-C2 | Biotechne |
| **CACNA1i** | Mouse | 459781-C1 | Biotechne |
| **PRKCG** | Mouse | 417911-C2 | Biotechne |
| **PRKCG** | Mouse | 417911-C1 | Biotechne |

**Table 2.**
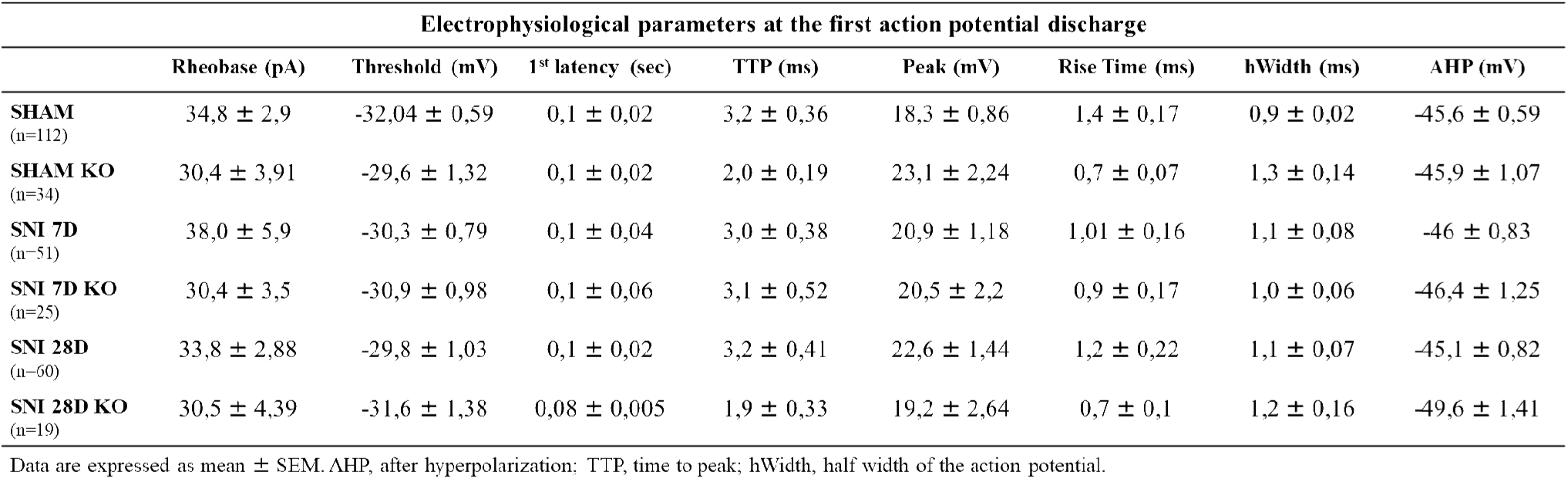
Electrophysiological parameters at the first action potential discharge.

### *In situ* hybridization

RNA was detected on 16 μm-thick sections using the RNAscope Multiplex Fluorescent Reagent Kit v2 (ACD, Biotechne, Minneapolis, MN) according, strictly, to manufacturer’s instruction. After tissue pre-treatment, sections were hybridized with mouse-specific probes (**Table 1**). Sections were counterstained with DAPI and mounted with ProLong Glass Antifade Mountant (Invitrogen™) for confocal imaging (Leica SP8-UV). Cell densities were normalized using the Cell Counter plugin of Fiji.

### Statistics

For cumulated distributions of amplitudes and intervals, bins were 0.5 pA and 0.05 s respectively. Not all these values were plotted for clarity, and a straight line represented the trajectories of the mean values. Otherwise noted, data were expressed as mean ± SEM. They were compared with the appropriate tests using GraphPad Prism (Boston, MA), and p<0.05 was considered as significantly different.

## Results

### Cav3.2 in PKCγ interneurons of the dorsal horn of the spinal cord

PKCγ interneurons are selectively examined in layer II-IV of the spinal cord using the PKCγ Cre ^ERT2^ mouse line x Ai14 (Abraira et al., 2017; Peirs et al., 2021). Immunofluorescence experiments performed on coronal slices of PKCγ Cre ERT2 x Ai14 mice showed that at least 50% of the neurons labelled with anti-PKCγ antibody were tdTomato positive (**Fig. 1A&B**). Ectopic expression of tdTomato was never found (**Fig. 1B**). PKC_ϒ_ neurons harbor Cav3.2 channels in various membranous compartments, including cell body, dendrites and synapses (**Fig. S1A, S1B**). Functional T-type calcium channels were recorded with the voltage-clamp configuration of the patch-clamp technique in PKCγ neurons detected by the live fluorescence of tdTomato (**Fig. 1C**). Both low-and high-threshold calcium currents were observed in current-voltage relationships in PKCγ neurons (**Fig. 1D**, **Fig. S1C**). On average, the current-voltage relationship of the low-threshold currents elicited from -100mV (n=16), were suppressed by Z944 (4 µM, n=5) a selective T-type calcium channel blocker (**Fig. 1E&F**). They were also partially inhibited by Ni^2+^ ions (90µM) by 43 ± 18 % of the control (n=10, p<0.001 vs control, Wilcoxon test, **Fig. 1F**). Since Cav3.2 has a much higher affinity for Ni^2+^ ions, as compared to Cav3.1 and Cav3.3, *in vitro*, (Lee et al., 1999), these data suggested that Cav3.2 participated in the low-threshold currents in PKCγ neurons, *in situ*.

**Figure 1.**
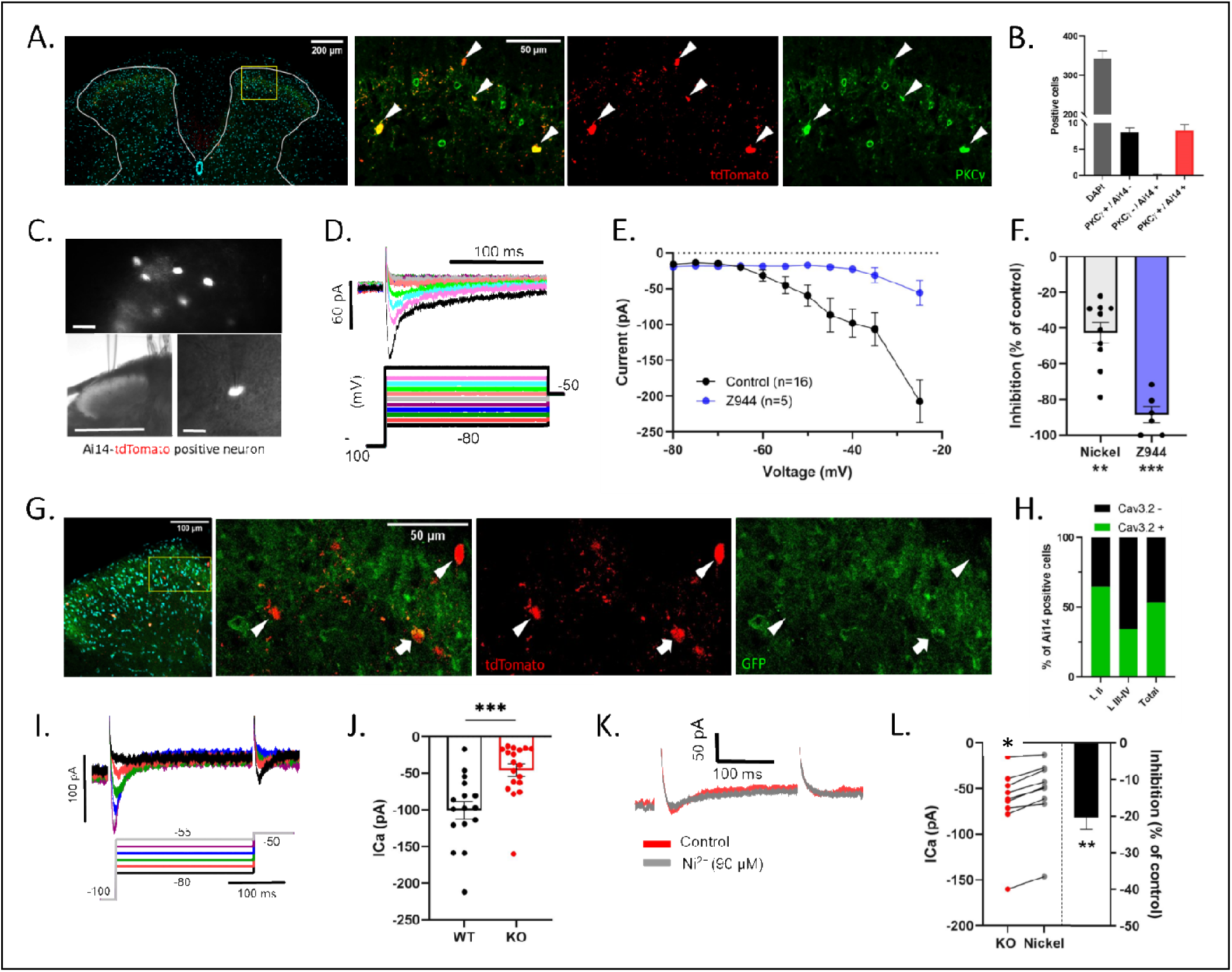
Identification of Cav3.2 in PKCγ interneurons. **A)** Immunostaining of a lumbar spinal cord slice (16 µm section) from adult PKCγ-Cre^ERT2^ x Ai14-tdTomato. This representative example shows the colocalization (arrowheads) of tdTomato (red) with PKCγ (green) in a DAPI (cyan) counterstained slice. **B)** Mean numbers of tdTomato and PKCγ positive neurons in 9 experiments similar as in **(A)** taken from 3 mice. **C)** Ai14-positive neurons (top) were resolved in acute lumbar slices from PKCγ-Cre^ERT2^ x Ai14-tdTomato expressing mice (40x magnification, scale bar 50 µm). They were selected for patch-clamp recording (bottom left, 4x magnification, scale bar 500 µm) using tdTomato as a marker (bottom right, 40x magnification, scale bar 25 µm). **D)** Family of typical T-type calcium currents elicited by increasing membrane potential, as indicated, from a holding potential of -100 mV, in a PKCγ neuron of a WT mice. **E)** Mean current-voltage relationship of T-type calcium currents in PKCγ neurons from WT mice (n=4) in the absence (n=16) and presence of Z944 (4 µM, n=5), recorded as in **(D)**. See Methods for details. Mean ± SEM are shown as symbols and lines. **F)** Summary of the inhibitory effects of Z944 (4 µM, n=6) and Ni^2+^ ions (90 µM, n=10) on the T-type calcium current of PKCγ neurons from WT mice (n=5). Mean ± SEM are shown as bars and lines. Dots show individual measurements. Different from control : ** p <0.01, *** p<0.001 Wilcoxon test. **G)** Immunodetection of GFP in a lumbar spinal cord slice (16 µm section) from adult PKCγ-Cre^ERT2^ x Ai14-tdTomato with a homozygous Cav3.2^GFP-Flox^ KI background. Arrowheads show tdTomato (red)-positive neurons lacking GFP labeling (green), and the arrow shows a colocalization of tdTomato with GFP. **H)** Proportions of Cav3.2 and Ai14 coexpressing neurons (n=265, 10 slices) in PKCγ-Cre^ERT2^ x Ai14-tdTomato x Cav3.2^GFP-Flox^ KI mice (n=2). Note that Cav3.2 and Ai14 coexpressing neurons were more numerous in lamina II as compared to lamina III-IV. **I)** Family of typical T-type calcium currents elicited at increasing membrane potentials as indicated, in a PKCγ neuron from a Cav3.2 KO mice. **J)** Comparison of T-type calcium current amplitudes in PKCγ neurons from KO (n=18, 6 mice) and WT (n=16, 5 mice) animals. *** p<0.001, Mann Whitney test. **K)** Example of the effect of Ni^2+^ 90µM on the T-type calcium current in a PKCγ neuron from a KO mice. **L)** Summary of the inhibitory effects of Ni^2+^ 90µM on the T-type calcium current of PKCγ neurons (n=9) of KO mice (n=5). Mean ± SEM are shown as bars and lines. Dots show individual measurements. Difference from control: ** p <0.01, Wilcoxon test.

The Cre recombinase-dependent ablation of cacna1h in PKCγ neurons was achieved in PKCγ Cre ERT2 x Ai14 mice with a homozygous Cav3.2^GFP-Flox^ KI background (referred as to KO in the above text), and compared to their wild-type (WT) counterpart. Immunofluorescence experiments aimed at verifying the level of Cre-dependent deletion of cacna1h in KO mice suggested that GFP labeling was excluded in only half of the PKCγ neurons where Cre-dependent expression of tdTomato was detected (**Fig. 1G&H**). However, the presence of Cav3.2 in PKCγ neurons was likely overestimated by GFP labeling because PKCγ neurons are located in an area enriched with GFP positive nerve terminals (François et al., 2015). Thus, the presence of Cav3.2 was verified at the functional level in patch-clamp experiments in KO mice (**Fig. 1I-L**). Low-threshold calcium currents were recorded in PKCγ neurons in KO mice (**Fig. 1I**), although their maximal amplitudes were smaller as compared to WT (-3.6 ± 0.6 pA/pF, n= 16 in WT and -1.6 ± 0.5 pA, n=18 in KO, p<0.0005, Wilcoxon, **Fig. 1J**). Furthermore, as shown in the typical example of **Fig. 1K**, the Ni^2+^ ions (90µM)-sensitive fraction of PKCγ neurons in KO mice (-22 ± 5 % of the control, n=8, p<0.001 vs control, Wilcoxon test) was lower than in WT mice (p<0.01 vs inhibition observed in WT, Mann Whitney test, **Fig. 1L and Fig.1F**). Altogether, these data established that a functional inactivation of Cav3.2 in PKCγ neurons was taking place in KO mice, accounting for some but not all the T-type calcium current in these neurons.

### Effects of Cav3.2 deletion in PKCγ interneurons on static mechanical sensitivity

Chronic pain is so far the major context requiring PKCγ neurons activity in rodents (Bardoni et al., 2013; Neumann et al., 2008; Peirs et al., 2015, 2021). Therefore, the role of Cav3.2 in PKCγ neurons was investigated during the course of allodynia in the spared-nerve-injury model (Decosterd & Woolf, 2000). A static mechanical allodynia was induced within days and remained stable throughout testing at the paw ipsilateral to the surgery in WT mice, unlike in sham animals (p<0.0001, 2-way ANOVA, **Fig. 2A**). KO mice had elevated static mechanical sensitivity before surgery (0.75 ± 0.08 g, n=16) as compared to WT (1.08 ± 0.09 g, n=17, p<0.05, Mann-Witney test, **Fig. 2A and C**) but the overall sensitization after surgery was similar in KO mice as compared to WT animals (p<0.001, 2-way ANOVA, **Fig. 2A**). However, Cav3.2 deletion in PKCγ neurons resulted in a transient decrease of pain seven days after surgery as compared to WT mice (respectively 0.036 ± 0.008 g and 0.013 ± 0.004 g, p<0.05, Mann-Witney test, **Fig. 2B**), which was no longer present at later time points. These behavioral data demonstrated a role for Cav3.2 in PKCγ neurons in the installation of allodynia.

**Figure 2.**
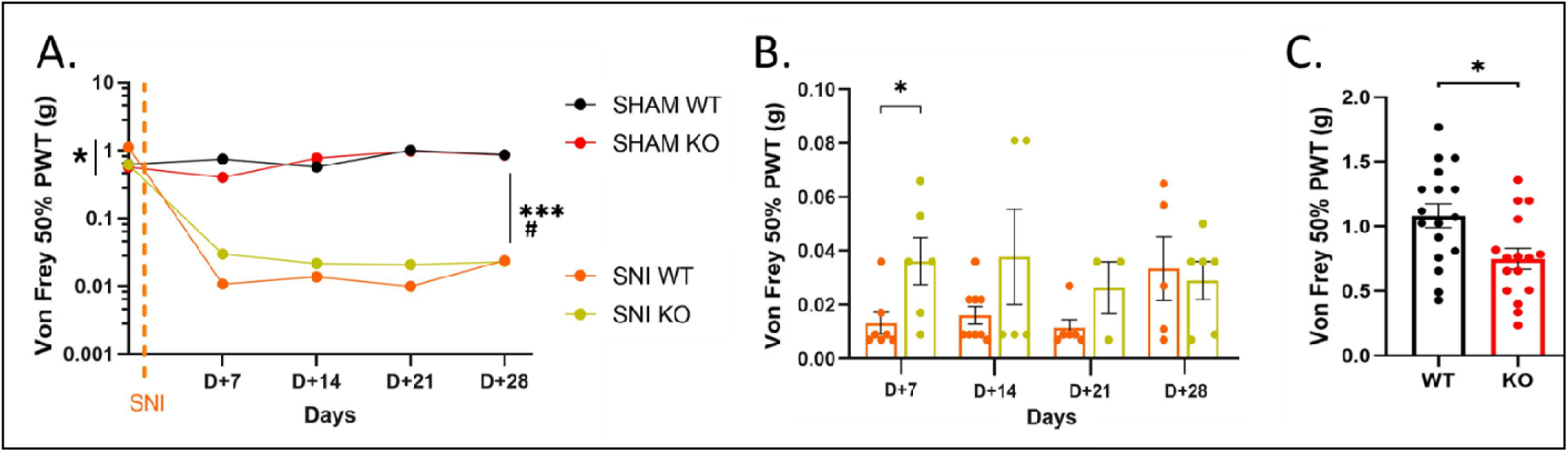
Early antalgic effect of Cav3.2 deletion in mice after Spared nerve injury. **A-C)** Static mechanical sensitivity was evaluated with the Von Frey filaments, using the up-and-down method, in PKCγ-CreERT2 x Ai14-tdTomato without (WT) and with a homozygous Cav3.2^GFP-Flox^ KI background (KO). **A)** Mean kinetics of the decreases of withdrawal thresholds after SNI at the ipsilateral paw in WT (n=9) and KO (n=7) mice (# for WT, p<0.0001 and *** for KO, p<0.001, 2-way ANOVA). Withdrawal thresholds were unaltered in WT (n=5) and KO (n=5) Sham littermates. **B)** An antalgic effect is seen 7 days after SNI in KO as compared to WT mice, and not during the later stages of the pathology. Same data as in **(A)**. **C)** Same as in **(B)** except that the comparison between WT (n=17) and KO (n=16) mice was performed before surgery. **(B, C)** Mean ± SEM are shown as bars and lines. Dots show individual measurements. *, p<0.05, Mann Whitney test.

### Roles of Cav3.2 in subthreshold properties in PKCγ neurons

T-type calcium channels participate in the intrinsic properties of some interneurons of the spinal cord (Harding et al., 2021; Ku & Schneider, 2011; Ruscheweyh & Sandkühler, 2002; Smith et al., 2015) and Cav3.2 deletion modifies action potential properties and firing patterns in the dorsal horn of the spinal cord (Candelas et al., 2019). Since the intrinsic electrophysiological properties of PKCγ neurons are not known, their subthreshold dynamics were first examined (after an hyperpolarizing conditioning prepulse at -90mV, Candelas et al., 2019; Dreyfus et al., 2010). In PKCγ neurons of Sham WT mice (n=111, **Fig. 3A**), this conditioning prepulse triggered rebound shaped depolarizations (96%), hyperpolarizing responses (2%) and flat responses (2%), as described in lamina II Cav3.2-positive interneurons (Candelas et al., 2019). These rebound potentials were also the most frequent responses in Sham KO mice (n=38, **Fig. 3A**) as well as in WT and KO mice after SNI (**Fig. S2A**). Data were identical in neurons from naive mice, as well as from Sham mice either 7 days or 28 days after surgery **(Fig. S2B).** Therefore, data from Sham mice were pooled in the present section. Rebound potentials of PKCγ neurons were potentiated by TTX 1µM as shown in the typical examples of **Fig. 3B** (Tringham et al., 2012). Indeed, the maximal activity of T-type channels is better resolved in the presence of TTX (because sodium channel threshold is more hyperpolarized than the potential for maximal activation of T-type calcium channels, Cain & Snutch, 2010). The maximal amplitude of the rebounds were similar in PKCγ neurons of Sham KO and WT mice in the presence (respectively 22 ± 3.6 mV, n=10, and 27 ± 1.6 n=33, **Fig. 3C**) and absence of TTX (**Fig. S2C**), suggesting that Cav3.2 was not mandatory under these conditions. Rebounds were also similar in SNI WT mice as compared to Sham WT mice in the absence of TTX (**Fig. S2D**). In marked contrast, when better resolved in the presence of TTX, the rebounds were significantly lower 7 days (21 ± 2.4 mV, n=17) and 28 days (20 ± 3.1 mV, n=14) after SNI as compared to Sham WT mice (both p<0.05, Mann Whitney test, **Fig. 3C**). This effect of SNI was no longer statistically significant in PKCγ neurons from KO mice (25 ± 3.8 mV, n=10 at 7 days; and 20 ± 3.8 mV, n=8 at 28 days, **Fig. 3C**). Note that the amplitude of the rebound was superimposable in males and females (**Fig. S2E**). The moderate effects of Cav3.2 ablation in PKCγ neurons was compared to the inhibition of the three T-type calcium channels with Z944 (1 µM). Superfusion with Z944 blunted the subthreshold rebound, as shown in the typical experiment of **Fig. 3E**. On average, this inhibitory effect of Z944 was similar in PKCγ neurons of WT mice, both in the absence (**Fig. S2G**) and in the presence of TTX (**Fig. 3F**). Z944 also inhibited the low-threshold rebound in KO mice, and this pharmacological effect reached significance in SNI mice, in the presence of TTX (p<0.05, Wilcoxon test, **Fig. 3F**) and not in the absence of TTX (when the resolution is not optimal, **Fig. S2G**). In conclusion, T-type calcium channels are responsible for the subthreshold rebounds in PKCγ neurons, where the role of Cav3.2 appeared undetectable in Sham mice, while unmasked after SNI.

**Figure 3.**
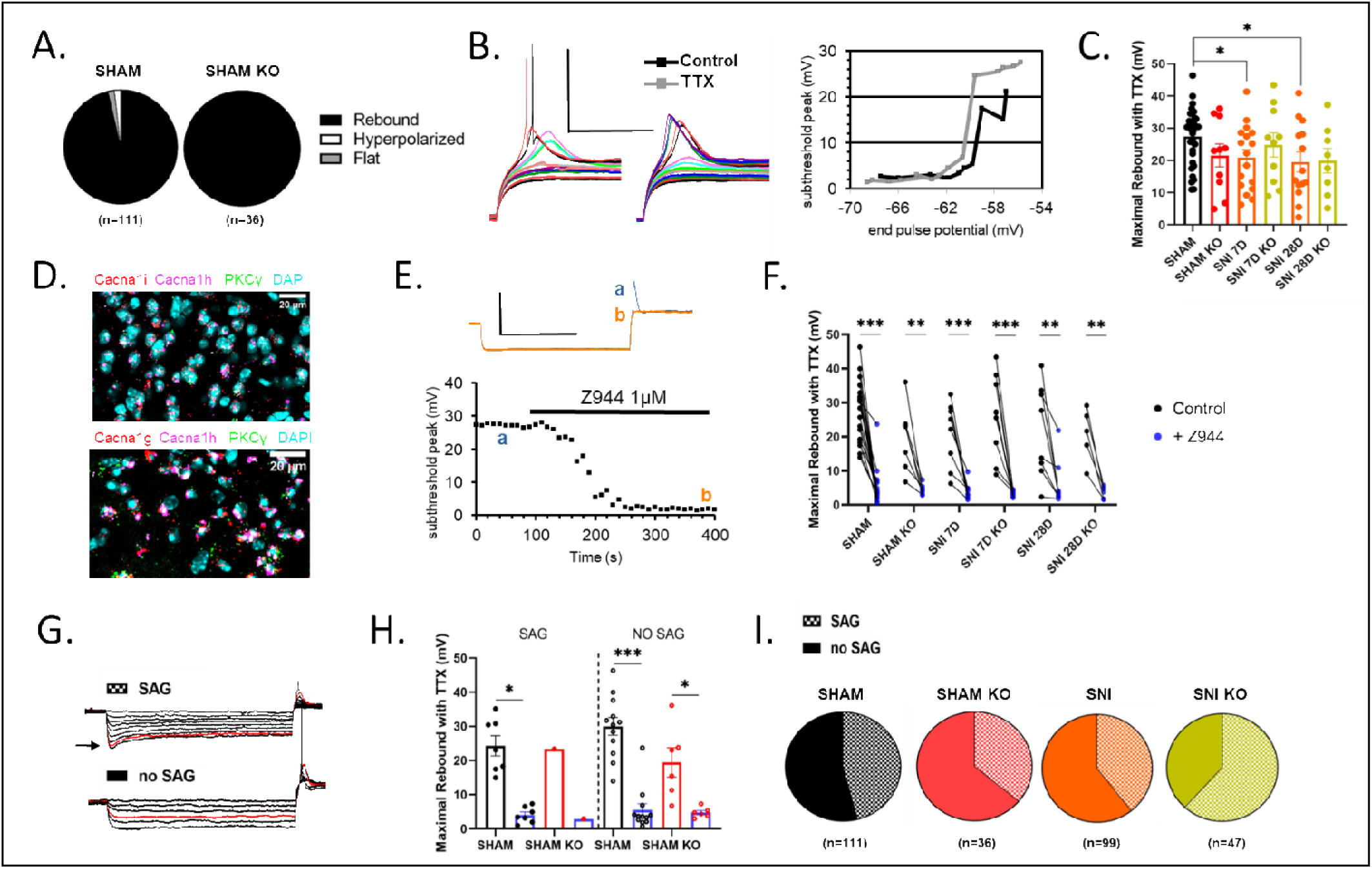
Involvement of T-type calcium channels in the subthreshold electrical properties of PKCγ neurons. **A)** Proportions of after-hyperpolarization responses (rebound, hyperpolarized, flat) recorded in PKCγ neurons of WT and KO Sham mice with the whole-cell patch clamp technique (see Methods for details). Rebounds were seen in 96% of the PKCγ neurons in WT mice. **B)** Representative membrane voltage traces recorded in the absence (left panel, Control) and presence of Tetrodotoxin (1 µM, TTX). The corresponding voltage-current relationship (right panel) shows the maximal amplitude of the subthreshold peak in the absence (black symbols) and presence of TTX (grey symbols). Holding current was -15pA; and increasing steps of 5pA were delivered from an initial level of 0pA. Bars are 50mV and 150ms. **C)** Maximal rebound amplitudes recorded in the presence of TTX (1 µM) in current-clamp experiments after SNI in WT and KO mice. Number of experiments were: 32 (Sham), 10 (Sham KO), 17 (SNI 7D), 10 (SNI 7D KO), 14 (SNI 28D), 8 (SNI 28D KO). Mean ± SEM are shown as bars and lines. Dots show individual measurements. *, p<0.05, Mann Whitney test. **D)** *In situ* hybridization directed against cacna1g (Cav3.1), cacna1h (Cav3.2), cacna1i (Cav3.3) and prkcg (PKCγ) in lumbar sections (16 µm-thick) of spinal cord from a PKCγ-CreERT2 x Ai14-tdTomato. **E)** Current-clamp experiment where the maximal amplitude of the subthreshold peak was recorded in the presence of TTX (1 µM) and in the presence of TTX + Z944 (1 µM) as indicated by the line. Representative voltage traces (top panel) were recorded at the time indicated by the corresponding letters on the main graph. Bars are 40mV and 1s. **F)** Effect of Z944 (1 µM) on maximal rebound amplitudes recorded in the presence of TTX (1 µM) in the current-clamp mode after SNI in WT and KO mice. Dots connected by a line are the data recorded from one neuron, before and during Z944 superfusion. Numbers of experiments were: 19 (Sham), 7 (Sham KO), 8 (SNI 7D), 9 (SNI 7D KO), 8 (SNI 28D), 5 (SNI 28D KO). **p < 0.01, ***p < 0.001, Wilcoxon test. **G)** Families of current-clamp recordings of a sag-and a no sag-exhibiting neurons. Note the presence of a transient deflection (arrow) during the 2-sec long hyperpolarisations in the sag-exhibiting neuron. Successive currents steps of decreasing values were applied from a holding of -70 mV. **H)** The mean effects of Z944 (1 µM) on the maximal rebound amplitudes were similar in sag-and no sag-neurons. The recordings were performed in the presence of TTX (1 µM) in WT and KO mice. Same experiments as in **(F)** ; Number were : 7 (sag-Sham), 1 (sag-Sham KO), 12 (no sag-Sham), 6 (no-sag Sham KO). Mean ± SEM are shown as bars and lines. Dots show individual measurements. *, p<0.05, ***, p<0.001, Wilcoxon test. **I)** The proportions of sag-(dotted areas) and no sag-PKCγ neurons (plain areas) in WT and KO mice after SNI are compared to those of Sham mice. Sag-neurons tended to be more numerous in SNI KO mice. Note that for SNI, data were pooled from the early and late time points.

The sag seen during the pre-pulse in some but not all recordings (**Fig. 3G**), suggested a role for hyperpolarized-activated channels (HCN). HCN channels were indeed reported recently in PKCγ neurons in transcriptomic databases and patch-clamp recordings in mice (Nakagawa et al., 2020). HCN can interfere with the activity of T-type calcium channels, in various ways including potentiation, compensation, or occlusion (Engbers et al., 2011, 2012; Hu, 1995). Sag-positive and sag-negative neurons exhibited similar residual rebound in the presence of Z-944 (∼5mV, **Fig. 3H**), suggesting that HCN channels were not strongly contributing to this subthreshold depolarization. Note that the fraction of sag-positive neurons tended to be larger in PKCγ neurons during SNI in KO mice (∼60%) as compared to WT and KO sham mice, and to WT SNI mice (∼40%) but this did not reach significance (**Fig. 3I**). Thus, HCN were not playing a differential role during pathology and/or in Cav3.2 deleted PKCγ neurons.

### Action potential properties of PKCγ neurons

In order to concentrate on the role of T-type channels during mechanical allodynia, we analyzed the action potential properties of the PKCγ neurons exhibiting a rebound potential (see above). The resting resistance was not changed after SNI, nor by Cav3.2 ablation (∼2.8GΩ, **Fig. 4A**), suggesting that a window current was not modified during allodynia, and was independent of Cav3.2. The rheobase was also similar among neurons (∼20 pA, **Fig. 4B**). At the rheobase, all the parameters of the first action potential were similar after SNI, whether Cav3.2 was present or not in PKCγ neurons (**Table 1**), including the latency to the first action potential (∼100 ms, **Fig. 4C**). Superfusion with Z944 revealed the role of T-type channels since the blocker diminished the resting resistance in PKCγ neurons of Sham mice (from 3 ± 0.6 to 1.9 ± 0.5 GΩ, n=14, p<0.01, Wilcoxon test, **Fig. S3A**). Z944 had a similar effect in Sham KO mice (from 4 ± 1.3 to 2.4 ± 0.5 GΩ, n=12, p<0.05, Wilcoxon test, **Fig. S3A**), and the effect was therefore independent of Cav3.2. Z944 also lowered the peak of the action potential in WT (from 19 ± 2.4 to 10 ± 2.4 mV, n=14, p<0.01, Wilcoxon test, **Fig. S3B**). Strikingly, Z944 did not change the peak of the action potential in KO mice, suggesting a cooperation of T-type channels, including Cav3.2, in sodium channel activation (**Fig. S3B**).

**Figure 4.**
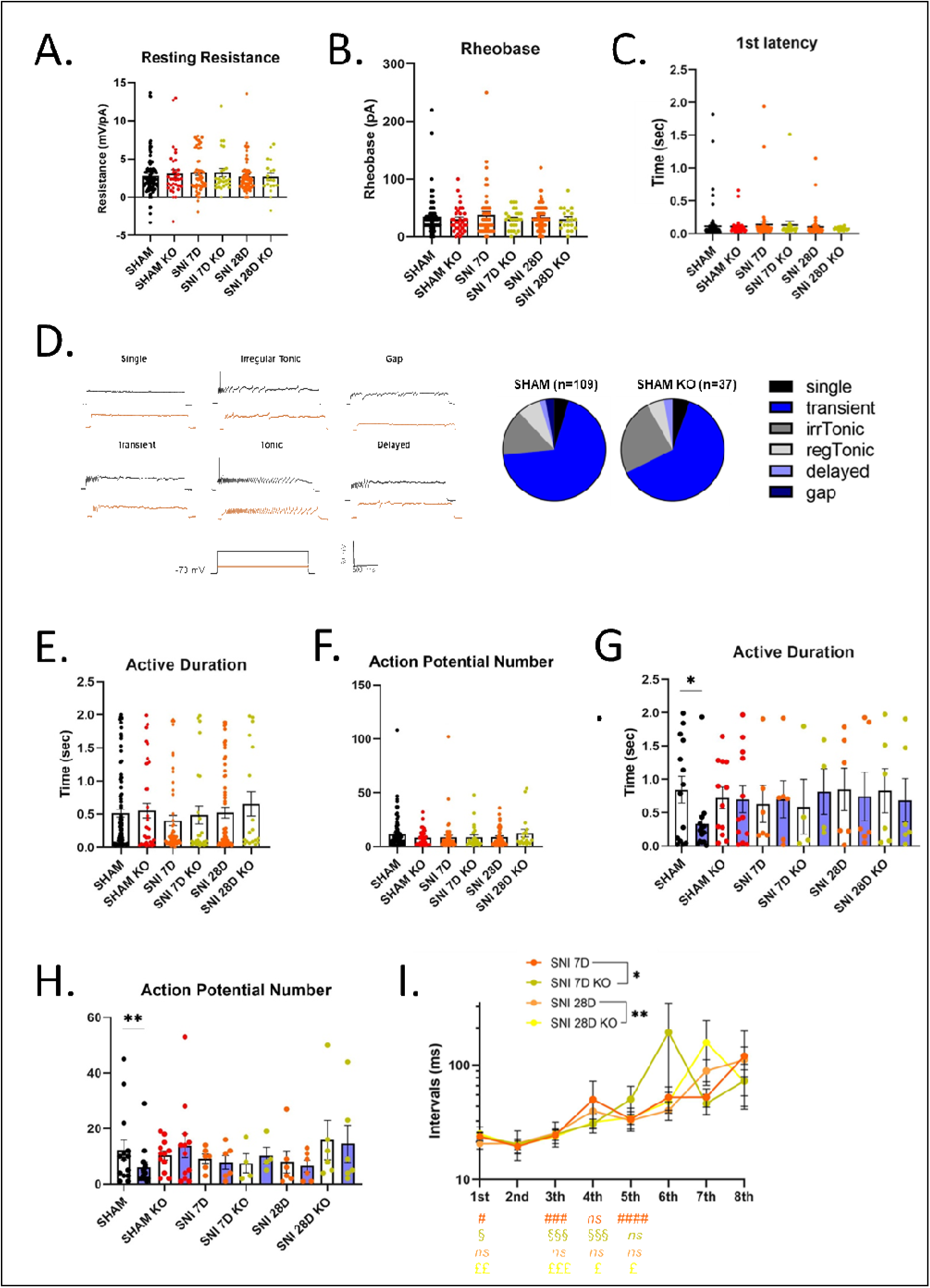
Role of T-type calcium channels in the excitability of PKCγ neurons. **(A-C)** SNI did not change the resting resistance (**A**), the rheobase (**B**) and the 1^st^ latency of the first action potential (**C**) elicited upon depolarisation from -70mV in PKCγ neurons of WT and KO mice. **(D)** Cav3.2 deletion did not modify the proportions of the firing patterns recorded in PKCγ neurons. **(E-F)** Effects of SNI on the active duration **(E)**, and the number of action potentials **(F)** during a 2s-long depolarisation in PKCγ neurons of WT and KO mice. **(G-H)** Active duration **(G)** and numbers of action potentials **(H)** of PKCγ neurons in the absence (white bars) and presence of Z944 (1 µM, blue bars) in WT and KO mice, after SNI. In **(A-C, E-F)** numbers of experiments were: 112 (Sham), 34 (Sham KO), 51 (SNI 7D), 25 (SNI 7D KO), 60 (SNI 28D), 19 (SNI 28D KO). Mean ± SEM are shown as bars and lines. Dots show individual measurements. *, p<0.05, Mann Whitney test. In **(G-H)** numbers of experiments were: 14 (Sham), 12 (Sham KO), 6 (SNI 7D), 4 (SNI 7D KO), 6 (SNI 28D), 6 (SNI 28D KO). Mean ± SEM are shown as bars and lines. Dots show individual measurements. *, p<0.05, ** p<0.01 Wilcoxon test. **(I)** The mean intervals of the first action potential during a 2s-long depolarisation in PKCγ neurons follow different relationships in WT and KO mice after SNI. Same experiments as in (**A**). Mean ± SEM are shown as symbols and lines. *, p<0.05, **, p<0.01, 2-way ANOVA. The lower panel summarized the statistical differences between the intervals of the 1st, 3rd, 4th or 5th intervals with the interval of the 2nd action potential. #, §, £ p<0.05 ; ££ p<0.01 ; ###, §§§, £££ p<0.001, #### p<0.0001, Wilcoxon test.

At maximal stimulation, the firing patterns of the Cav3.2-positive interneurons of the lamina II of the spinal cord are heterogeneous (Candelas et al., 2019). In PKCγ neurons of both WT and KO Sham mice, ∼90% of the firing patterns were brief (single, transient and irregular tonic, **Fig. 4D**). This distribution of firing patterns was unaltered after SNI (**Fig. S3C**), which never showed lengthening of the activity. Furthermore, the activity duration, and the number of action potentials, triggered during the 2-s pulse, were unchanged after SNI (**Fig. 4E&F**). Cav3.2 deletion did not change these parameters (**Fig. 4E&F**) but the non-selective inhibition of T-type channels by Z-944 halved the activity duration (from 0.84 ± 0.2 s to 0.33 ± 0.1 s, n=14, p<0.05, Wilcoxon test) and diminished the number of action potentials (from 12 ± 4 to 6 ± 2, n=14, p<0.01, Wilcoxon test) in WT Sham mice (**Fig. 4G&H**). This effect was absent in KO Sham mice, where Z-944 no longer change the activity duration (from 0.72 ± 0.2 s to 0.70 ± 0.2 s, n=12) and the number of action potentials (from 10 ± 1.6 to 14 ± 4.3, n=12, **Fig. 4G&H**). These firing properties were profoundly modified after SNI (**Fig. 4G&H**) since activity duration and the number of action potentials were not modulated by either or both Z-944 and Cav3.2 deletion. Thus, PKCγ neurons undergo a remodeling after SNI: while the firing pattern is T-type dependent and involves Cav3.2 in Sham mice, it seems dominated by ion channels other than T-type calcium channels in a Cav3.2-insensitive manner during the pathology.

The firing patterns were examined further in PKCγ neurons after SNI, since two studies reported a pairing between the first and the second action potential in Cav3.2-expressing neurons (Candelas et al., 2019; Dumenieu et al., 2018). The interval of the 2^nd^ action potential was indeed the shortest of the first action potentials in PKCγ neurons in both Sham WT and Sham KO mice (Wilcoxon test, **Fig. S3D**). The overall U-shape of these relationships was superimposable in Sham WT and Sham KO mice (**Fig. S3D**). Seven days after SNI, the U-shape of the mean distribution was flattened in WT mice (n=29), but not in KO mice (n=15, p<0.05, 2-way ANOVA, **Fig. 4I**). This difference between WT and KO mice was maintained at the later time point of the pathology (respectively n=37 and n=15, p<0.01, 2-way ANOVA, **Fig. 4I**). Differences between the interval of the 2^nd^ action potential and the others were still present 7 days after SNI, but absent 28 days after SNI in WT mice (**Fig. 4I**). The U-shape distribution was restored in KO mice, 28 days after SNI (**Fig. 4I**). Thus, an involvement of Cav3.2 in fine patterning of the firing of PKCγ neurons, undetectable in Sham mice, was potentiated 7 days after SNI and further increased during the late stage of the model.

### Effects of Cav3.2 deletion in PKCγ neurons on inhibitory neurotransmission

The inhibitory control of PKCγ neurons is impaired in mice models of chronic pain exhibiting mechanical allodynia (Bardoni et al., 2013; Sandkühler, 2009). Since T-type calcium channels including Cav3.2 modifies inhibitory synapse activity (Candelas et al., 2019; Leresche & Lambert, 2017), the IPSC of PKCγ neurons were recorded in acute lumbar spinal cord slices from adult mice using the whole cell configuration of the patch-clamp technique held at a membrane potential of -30mV (**Fig. 5A,B**) (Candelas et al., 2019). Amplitudes and inter-events intervals of the spontaneous IPSC were indistinguishable in PKCγ neurons from sham mice early (7 days) and late (28 days) after surgery in either WT (Sham 7 days n=69; Sham 28 days n=63) or KO backgrounds (Sham 7 days n=24, Sham 28 days n=16, **Fig. S4A**). IPSCs were similar whether PKCγ neurons were located in the dorso-ventral axis (lamina II to IV, supplementary **Fig. S4B**) and in the coronal axis of the lumbar spinal cord (subdivided in 3 areas, supplementary **Fig. S4C**).

**Figure 5.**
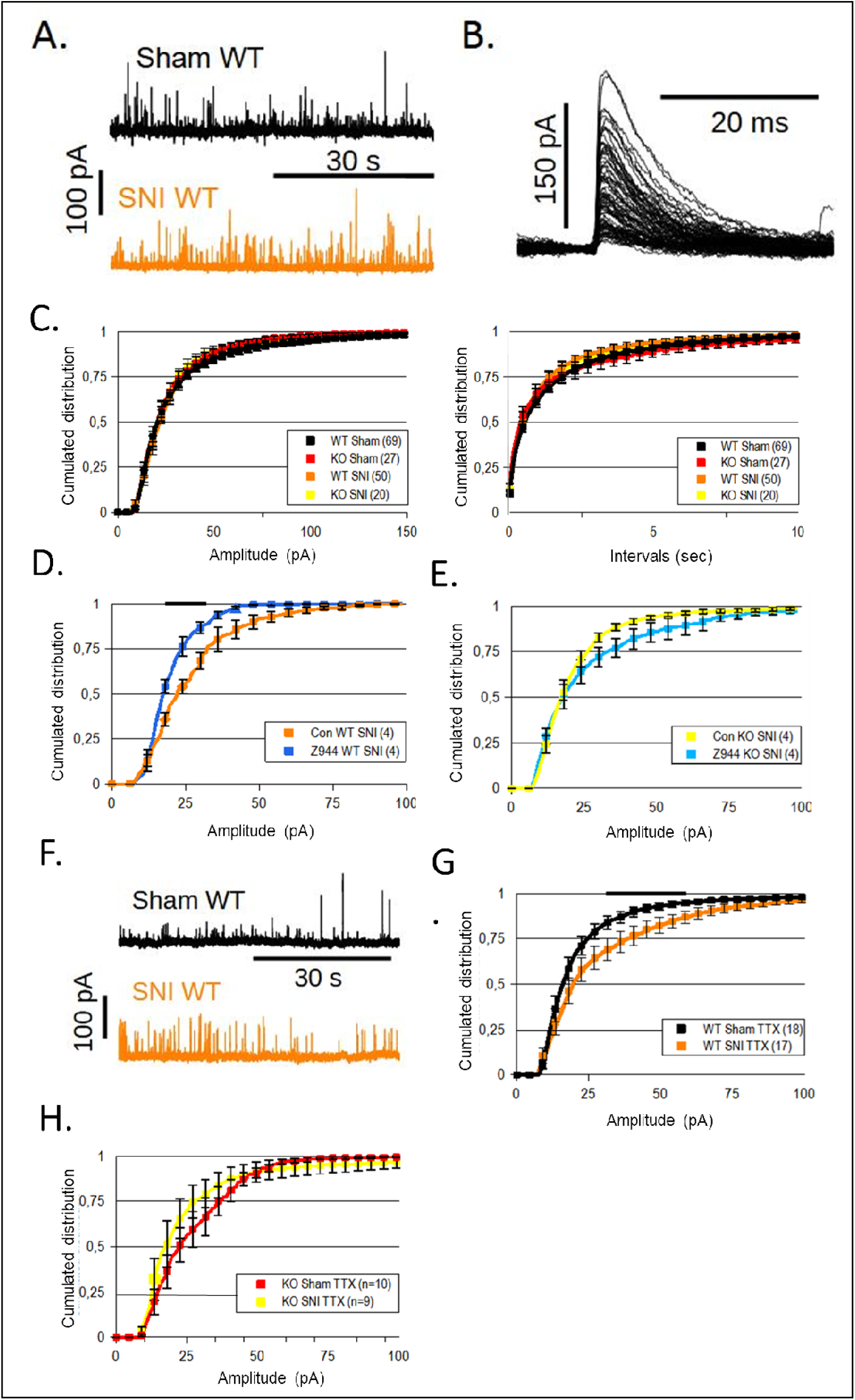
Early changes in the inhibitory neurotransmission of PKCγ neurons. **(A-E)** Inhibitory synaptic currents recorded 7 days after SNI. **(A)** Whole-cell voltage clamp recordings of ISPC of PKCγ neurons from WT Sham and SNI mice. **(B)** Family of ISPC extracted from a recording as in (**A**), in a WT Sham mince. (**C**) IPSC amplitudes (left panel) and intervals (right panel) were unchanged in PKCγ neurons in either WT of KO mice, early after SNI. **(D,E)** The T-type channel blocker, Z944 (1 µM) diminished IPSC amplitudes early after SNI in WT (**D**) and not in KO mice **(E). (F-H)** Miniature IPSC recorded in the presence of the sodium channel blocker, TTX (1 µM). **(F)** Raw traces of mIPSC of PKCγ neurons from WT Sham and SNI. **(G,H)** mIPSC were larger early after SNI in WT mice **(G)** and not in KO mice **(H)**, as compared to their Sham counterparts. Graphs are cumulated distributions of the number of recordings indicated within brackets. Mean ± SEM are shown as symbols and lines. When present, a black bar indicates the range of values where statistical differences between appropriate distributions were larger than 0.05, using a Wilcoxon text **(D,E)** or using a Mann Whitney test (**C,G,H**).

At an early time point (7 days), IPSC were similar in sham and SNI mice (n=69 and 50 and respectively, **Fig. 5C**) and the ablation of Cav3.2 in KO mice, did not modify this comparison (n=27 and 20 respectively, **Fig. 5C**). IPSCs were not significantly modified by Z-944 in Sham WT and KO mice (n=9 and 7 respectively, **Fig. S4D&E**), ruling out a role for T-type channels under physiological conditions. In contrast, Z-944 significantly reduced the amplitudes by ∼20% of the IPSC in PKCγ neurons of SNI WT mice (n=4, p<0.05, Wilcoxon test, **Fig. 5D**), and this effect was blunted in SNI KO mice (n=4, **Fig. 5E**). Altogether, these results showed that SNI potentiated the roles of T-type calcium channels in the inhibitory network of PKCγ neurons, and that Cav3.2 deletion in PKCγ neurons blunted this remodeling., Miniature IPSC (mIPSC) were next examined in PKCγ neurons in the presence of TTX, the sodium channel blocker which isolates synapses from the influence of the network (**Fig. 5F**). The amplitudes of mIPSC were larger in SNI WT mice as compared to their sham siblings (n=17 and 18 respectively, p<0.05, Mann Whitney test, **Fig. 5G**) and this change was not seen in KO mice (sham n=17, SNI n=9, ns, Mann Whitney test, **Fig. 5H**). Thus, Cav3.2 ablation in PKCγ neurons prevented from the remodeling induced by SNI of some inhibitory synapses of PKCγ neurons.

At a late stage of the disease (28 days), the amplitude of the IPSC were lower in SNI mice as compared to sham mice with a WT background (n=28 and 63 respectively, p<0.05, Mann Whitney test, **Fig. 6A,B**), but not with a KO background (sham n=17, SNI n=18, ns, Mann Whitney test, **Fig. 6C**). Importantly, Z944 did not modify either the amplitude or the intervals of the IPSC in any groups (**Fig. S4F&G**) suggesting that 1) T-type channels were not mandatory for IPSC in sham mice, and 2) at least two T-type channels, including Cav3.2 of PKCγ neurons, were involved in the inhibitory network of PKCγ neurons, 28 days after SNI. The distributions of the mIPSC amplitudes were similar in Sham and SNI mice (**Fig. 6D,E**). However, the intervals of the mIPSC were lower in the PKCγ neurons of the SNI KO mice as compared to the SNI WT mice (n=7 and 20 respectively, p<0.05 Mann Whitney test, **Fig. 6F**), unlike in Sham WT and KO mice. Thus, the effects of SNI on the inhibitory transmission were absent after Cav3.2 deletion in PKCγ neurons, and the activity of Cav3.2 at some inhibitory synapses of PKCγ neurons were detectable during SNI only.

**Figure 6.**
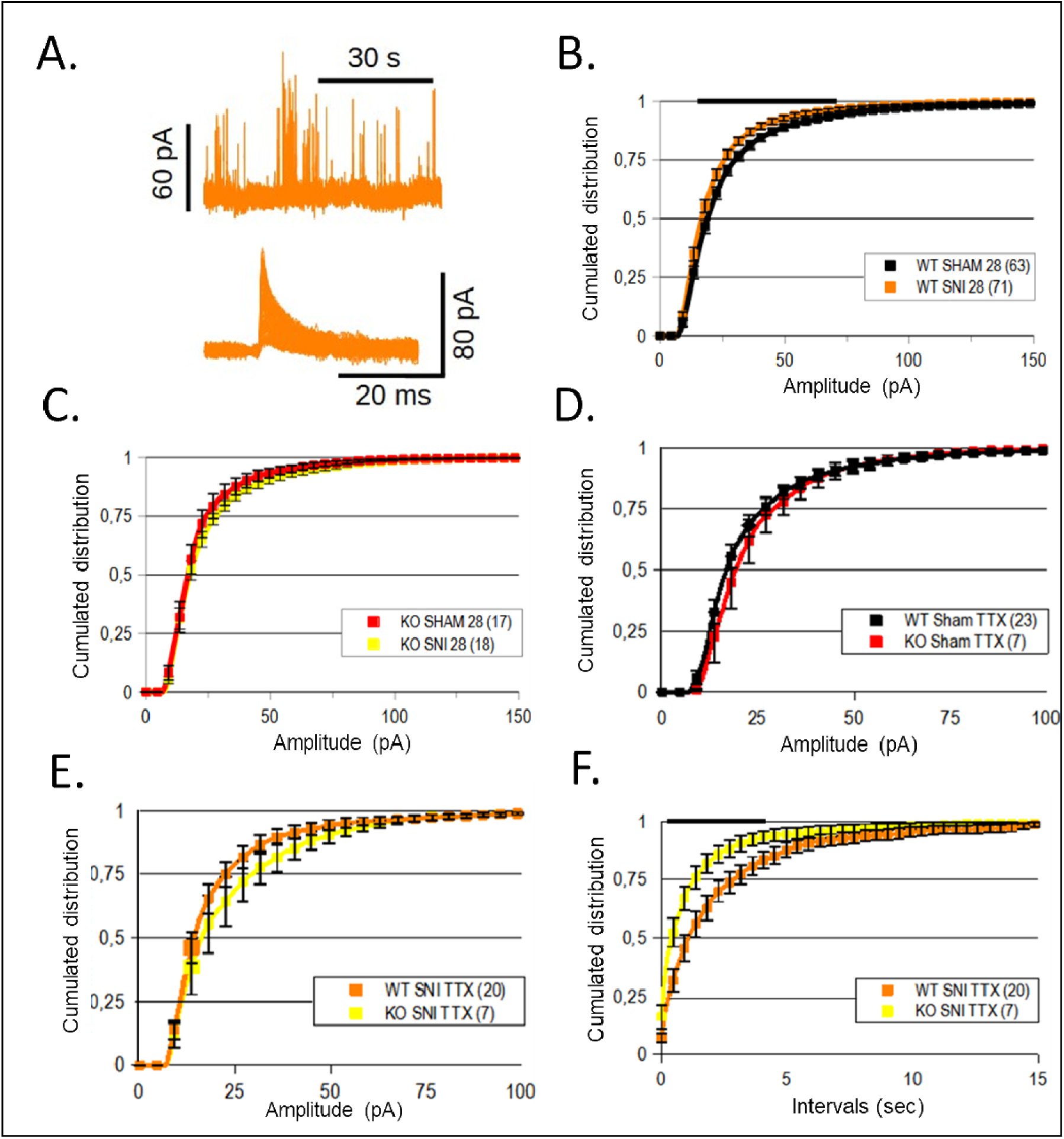
Late changes in the inhibitory neurotransmission of PKCγ neurons. **(A-C)** Inhibitory synaptic currents recorded 28 days after SNI. **(A)** Whole-cell voltage clamp recording of ISPC of PKCγ neurons from a SNI mice (top panel) and a family of ISPC extracted from this recording (bottom panel). **(B,C)** IPSC amplitudes were smaller in PKCγ neurons of WT mice **(B)**, and not in KO mice **(C)** late after SNI. **(D,E)** Miniature IPSC amplitudes were similar in PKCγ neurons of Sham mice **(D)**, and in SNI mice **(E)** late after SNI. **(F)** Miniature IPSC intervals were smaller in PKCγ neurons of KO mice as compared to WT mice, late after SNI. Graphs are cumulated distributions of the number of recordings indicated within brackets. Mean ± SEM are shown as symbols and lines. When present, a black bar indicates the range of values where statistical differences between appropriate distributions were larger than 0.05, using a Mann Whitney test (**B-F**).

### Effects of Cav3.2 deletion in PKCγ neurons on excitatory neurotransmission

Remodeling of the glutamatergic neurotransmission is taking place in the spinal cord during normal and pathological pain processing (Inquimbert et al., 2012; Jacus et al., 2012; Kopach et al., 2015; Sandkühler, 2009). T-type calcium channels including Cav3.2 are well-known to participate in glutamatergic synaptic transmission (Isope et al., 2012; G. Wang et al., 2015). The spontaneous excitatory synaptic currents (EPSC) of PKCγ neurons were therefore recorded in the lumbar slices at a membrane potential of -70mV (**Fig. 7A,B**) (Candelas et al., 2019). The properties of these EPSC did not change in PKCγ neurons between the early time point (7 days) and the later time point (28 days) in sham WT (7 days n=53, 28 days n=54) and KO mice (7 days n=21, 28 days n=17, **Fig. S5A**). EPSCs were similar whether PKCγ neurons were located in either axis of the coronal slices (**Fig. S5B&C**).

**Figure 7.**
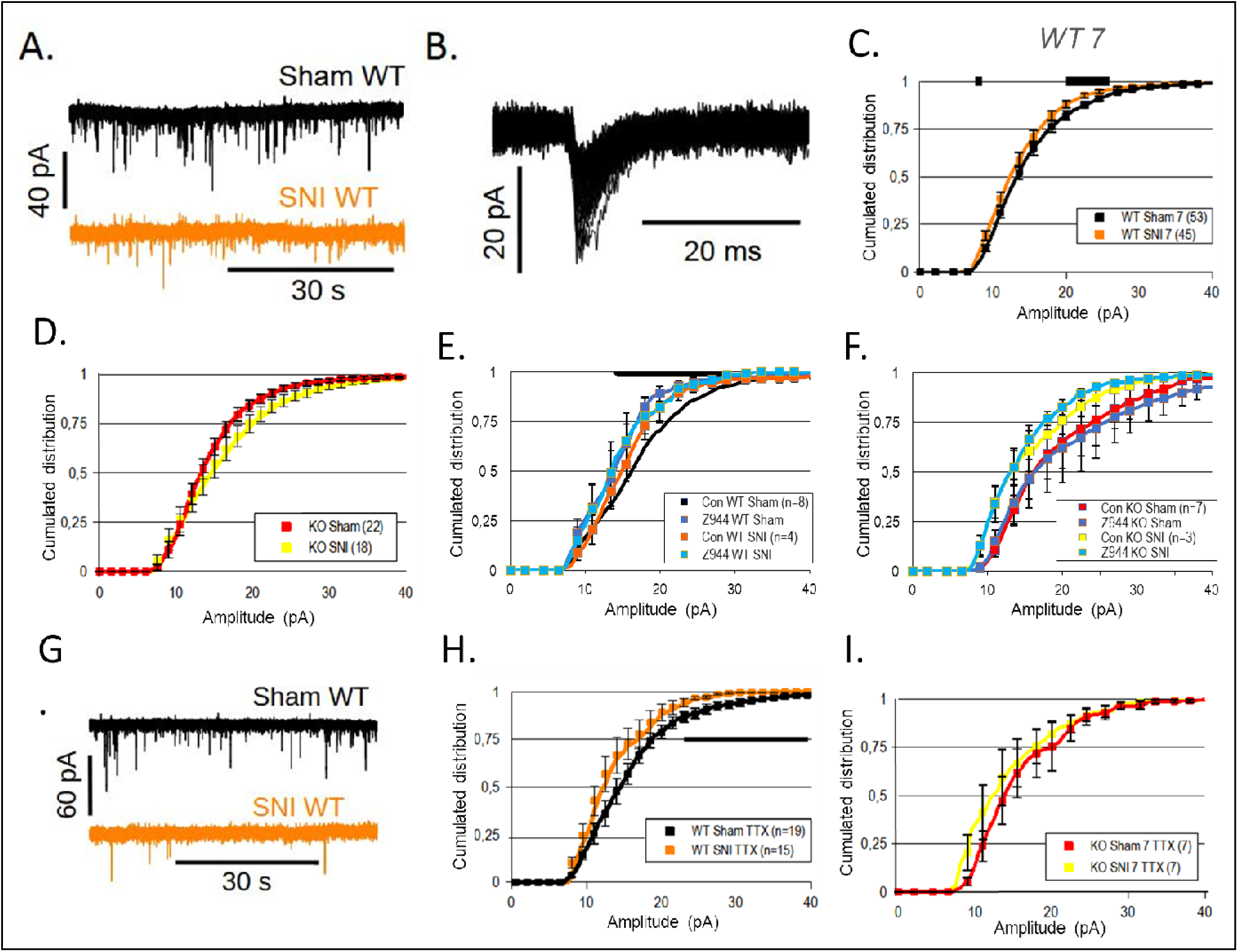
Early changes in the excitatory neurotransmission of PKCγ neurons after SNI. **(A-F)** Excitatory synaptic currents recorded 7 days after SNI. **(A)** Whole-cell voltage clamp recordings of ESPC of PKCγ neurons from WT Sham and SNI mice. **(B)** Family of ESPC extracted from a recording as in (A), in a WT Sham mice. **(C)** EPSC amplitudes were smaller in PKCγ neurons of WT mice, 7 days after SNI, as compared to WT Sham mice. and were superimposable in PKCγ neurons of Sham and SNI KO mice **(D)**. **(E,F)** The T-type channel blocker, Z944 (1 µM) diminished EPSC amplitudes in Sham WT mice **(E)** and not in Sham KO mice **(F)**, while Z944 did not change the amplitude in WT and KO mice, 7 days after SNI **(E,F). (G-I)** Miniature EPSC recorded in the presence of the sodium channel blocker, TTX (1 µM). **(G)** Raw traces of mEPSC of PKCγ neurons from WT Sham and SNI. **(H,I)** mIPSC were smaller 7 days after SNI in WT mice **(H)** and not in KO mice **(I)**. Graphs are cumulated distributions of the number of recordings indicated within brackets. Mean ± SEM are shown as symbols and lines. When present, a black bar indicates the range of values where p<0.05 between appropriate distributions were larger than 0.05, Wilcoxon test **(E,F)** and Mann Whitney test (other graphs).

The amplitudes of the EPSC were diminished early during SNI in WT mice as compared to the sham WT mice (n=45 and 53 respectively, p<0.05, Mann Whitney test, **Fig. 7C**), and this difference was not observed in KO mice (Sham n=22 and SNI n=18, ns, Mann Whitney test, **Fig. 7D**), accounting for the gap between EPSC distributions in SNI KO as compared to SNI WT mice (p<0.05, Mann Whitney test, **Fig. S5D**). Thus, Cav3.2 of PKCγ neurons is mandatory for a moderating effect of SNI on excitatory transmission. In addition, the T-type blocker Z944 diminished the EPSC in PKCγ neurons of sham WT mice (n=8, significant in 50% of the events at p<0.05 Wilcoxon test, **Fig. 7E**) and not in PKCγ neurons of KO sham (n=7, **Fig. 7F**). Z944 had no significant effects in PKCγ neurons of SNI WT (n=4, **Fig. 7E**) and KO (n=3, **Fig. 7F**) mice. Altogether, these findings suggested that Cav3.2 channels were participating in EPSC of PKCγ neurons in Sham mice, an effect that was no longer active in SNI animals. In order to focus on excitatory synapses of PKCγ neurons, miniature EPSC (mEPSC) were recorded in the presence of TTX (**Fig. 7G**). The amplitudes of mEPSC were smaller in PKCγ neurons of WT SNI mice as compared to WT sham mice (respectively, n=15 and 19, p<0.05, Mann Whitney test, **Fig. 7H**) and this difference was absent in PKCγ neurons of KO mice (Sham n=7, SNI n=7, **Fig. 7I**). Thus, Cav3.2 is moderating some glutamatergic synapses of PKCγ neurons during SNI, and its post-synaptic deletion normalized the synaptic activity early after SNI.

EPSC were examined 28 days after surgery (**Fig. 8A**). First, Z944 diminished EPSC in PKCγ neurons of sham WT mice (n=7, significant for ∼12% of the events p<0.05 Wilcoxon test, **Fig. 8B**), and this effect was weaker than 7 days after SNI (see **Fig. 7E**). In addition, the amplitudes of the spontaneous EPSC and of the mEPSC were smaller in PKCγ neurons of WT SNI (EPSC, n=58; mEPSC, n=15) as compared to WT sham (EPSC, n=54; mEPSC, n=20, p<0.05, Mann Whitney test for both comparisons, **Fig. 8C&E**), again showing that mechanical allodynia was moderating the glutamatergic network of PKCγ neurons. In contrast, spontaneous EPSC and mEPSC were superimposable in Sham (EPSC n=18; mEPSC n=28) and SNI (EPSC n=14; mEPSC n=4) KO mice (**Fig. 8D&F**), again suggesting that Cav3.2 of PKCγ neurons was mandatory in this remodeling of the excitatory synapses of PKCγ neurons The excitatory neurotransmission was therefore not different at 28 and 7 days after SNI.

**Figure 8.**
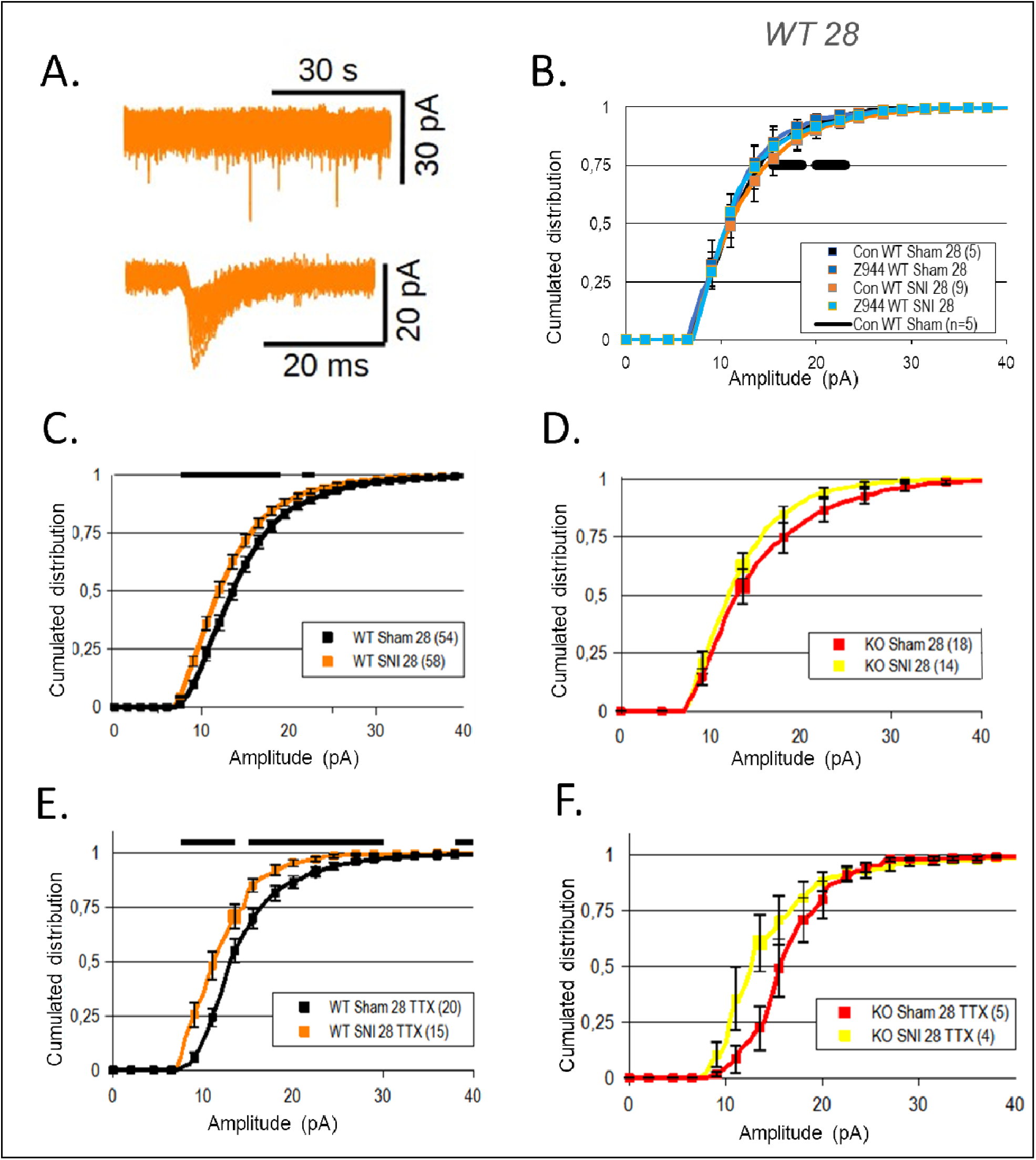
Late changes in the excitatory neurotransmission of PKCγ neurons after SNI. **(A-D)** Excitatory synaptic currents recorded 28 days after SNI. **(A)** Whole-cell voltage clamp recording of ESPC of PKCγ neurons from a SNI mice (top panel) and a family of ESPC extracted from this recording (bottom panel). **(B)** The T-type channel blocker, Z944 (1 µM) EPSC amplitudes in Sham WT mice and not in Sham WT SNI**. (C,D)** EPSC amplitudes were smaller in PKCγ neurons of WT mice **(C)** and not in KO mice **(D)** 28 days after SNI. **(E)** Miniature EPSC amplitudes were smaller in PKCγ neurons of WT mice, and not in KO mice **(F)** 28 days after SNI. Graphs are cumulated distributions of the number of recordings indicated within brackets. Mean ± SEM are shown as symbols and lines. When present, a black bar indicates the range of values where p<0.05 between appropriate distributions were larger than 0.05, Wilcoxon test **(B)** and Mann Whitney test **(C-F)**.

## DISCUSSION

The Cav3.2^GFP-Flox^ KI mouse was instrumental in deciphering the role of Cav3.2 in primary afferent neurons during chronic pain (François et al., 2015), in dorsal horn interneurons of the spinal cord (Candelas et al., 2019), and in parvalbumin-expressing neurons in the thalamus (Fayad et al., 2022). This mouse genetic model was used here for ablating Cav3.2 in PKCγ neurons of the spinal cord and examining the role(s) of this T-type calcium channel subtype during the development of chronic pain after spared-nerve injury (Decosterd & Woolf, 2000). This approach targeted a critical neuronal subtype in the gate control circuits of pain within the spinal cord, which can, during mechanical allodynia, transmit excitations from low-threshold mechanical pathways to high-threshold nociceptive projection neurons (Bardoni et al., 2013; Lu et al., 2013; Peirs et al., 2021; Q. Wang et al., 2020). Interestingly, SNI induced a remodeling of the electrical and synaptic activities of PKCγ neurons in WT mice, and these changes were either blunted or absent in KO mice.

There are evidences for the presence of several T-type calcium channels in PKCγ neurons of the spinal cord (Candelas et al., 2019; Russ et al., 2021, the present study). The inhibitory effect of a moderate concentration of Ni^2+^ ions suggested that Cav3.2 accounted for a maximum of ∼40% of the T-type calcium current, like in Y1 receptor-expressing interneurons (Sinha et al., 2021) and less than in some calretinin-positive interneurons of the dorsal horn of the spinal cord (Smith et al., 2015). In Cav3.2-expressing interneurons of Lamina II, Cav3.1 and Cav3.2 were not redundant since the subthreshold and suprathreshold properties of Cav3.2-expressing neurons were similar in Cav3.1 knockout mice and in wild-type mice (Candelas et al., 2019). In contrast, the sole Cav3.2 ablation in PKCγ neurons tended to lower the subthreshold rebound, although this was not significant, and did not change parameters influenced by T-type calcium channels in neurons (resting membrane potential, rheobase, action potential latency) (see reports by Cain & Snutch, 2010; Candelas et al., 2019; Deleuze et al., 2012; R. Wang & Lewin, 2011). This was unexpected since Cav3.2 ablation in PKCγ neurons had strongly lowered the amplitude of T-type calcium currents, as well as the fraction of Ni^2+^-sensitive T-type current. Of note, others have reported that T-type calcium current amplitudes are not predictive of their involvement in the excitability of primary afferent neurons (R. Wang & Lewin, 2011), and thalamic neurons (Deleuze et al., 2012). The weak coupling between Cav3.2 and action potential initiation might be related to the absence of Cav3.2 in the axon initial segment of PKCγ neurons, unlike in cortical neurons (Lipkin et al., 2021) and granular neurons of the hippocampus (Dumenieu et al., 2018). These findings also suggested that Cav3.2 was not mandatory for firing in the 5-10 % of PKCγ neurons classified as “single” firing, which conveys spike time precision and the opportunity in pairing synaptic inputs on postsynaptic neurons, on a millisecond time scale (Ku & Schneider, 2011; Prescott & Koninck, 2002; Ratté et al., 2015). Nevertheless, the resting resistance was modified and the subthreshold rebound was inhibited by Z944. T-type channel(s) other than Cav3.2 appeared important in the modulation of the low-threshold spike of PKCγ neurons (Cain & Snutch, 2010; Liao et al., 2011; Llinás & Steriade, 2006). Z-944 did not modify the properties of the action potential, either, despite a large inhibitory effect on the rebound potential. This is in contrast to previous findings where T-type calcium channels, including Cav3.2, were found important in single spiking, especially in neurons expressing low levels of T-type calcium channels (Cain & Snutch, 2010; Candelas et al., 2019). However, in another study of unidentified neurons of the dorsal spinal cord, transient firing patterns were simply governed by sodium channel availability (Melnick et al., 2004), a mechanism that might take place in PKCγ neurons.

The amplitude of the action potential, the active duration and the number of action potentials were not modified by Z-944 perfusion after Cav3.2 deletion, showing that Cav3.2 was necessary but not sufficient in modulating firing (in PKCγ neurons classified as “not single”, though). Neither Cav3.2 ablation, nor Z-944 perfusion did change the relative distribution of firing patterns, although T-type calcium channels support transient activities when present in large amounts, or tonic activity when present in lower amounts (Cain & Snutch, 2010; Llinás & Steriade, 2006). The action potential pairing, eliminated by Cav3.2 ablation in Cav3.2-expressing interneurons of lamina II (Candelas et al., 2019) and in mature granule neurons of the dentate gyrus (Dumenieu et al., 2018), was not changed in PKCγ neurons of Sham KO mice as compared to WT animals, again supporting the view that Cav3.2 was not mandatory for the intrinsic electrical activity of PKCγ neurons.

SNI is known to modify the electrical activity of PKCγ neurons evoked by stimulations of primary afferent neurons, but little is known about the intrinsic properties of these neurons (Abraira et al., 2017; Lu et al., 2013; Peirs et al., 2021; Q. Wang et al., 2020; Zhi et al., 2022). The intrinsic properties of inhibitory neurons are unchanged after sciatic nerve constriction (CCI model, (Gassner et al., 2013; Schoffnegger et al., 2006), while in the SNI model, they are decreased in parvalbumin neurons (Boyle et al., 2019), and facilitated in Y1R-expressing neurons (Nelson et al., 2022). SNI lowered the subthreshold rebound amplitude without changing the action potential properties and the relative proportions of firing patterns of PKCγ neurons. Fine spike tuning was also modified after SNI since the U-shape distribution of the action potentials was flattened, and the pairing between the first action potentials was absent 28 days after SNI. The above changes occurring after SNI were not observed in Cav3.2-ablated PKCγ neurons. Thus, 1) SNI exacerbated a role for Cav3.2 in action potential firing of PKCγ neurons (**Fig. 9A,B**), and 2) Cav3.2 ablation restored Sham levels. Importantly, Z-944 no longer modified the gross firing properties of PKCγ neurons at the early and late stages after SNI, and this was not normalized by Cav3.2 ablation. Multiple ionic channels are modified during mechanical allodynia in the dorsal horn of the spinal cord, including sodium and potassium channels (Chen et al., 2009; Dou et al., 2018; Garcia et al., 2019; Ippolito et al., 2005; Sun et al., 2002; Yoo et al., s. d.), and they might play a role in molecular remodelling of PKCγ neurons during mechanical allodynia. Regarding the gate control mechanism, while most PKCγ neurons had a “brief” activity (i.e., single + transient + irregular tonic), 80% of the GlyT2-iCre-tdTomato neurons inhibiting PKCγ neurons are tonic (He et al., 2021; Ma et al., 2023). The distribution patterns of PKCγ neurons were not modified after SNI while the inhibitory (parvalbumin) neurons are turned from tonic to transient (Ma et al., 2023). Therefore, it will be interesting to characterize the firing pattern of PKCγ neurons during activation of primary afferent neurons.

**Figure 9.**
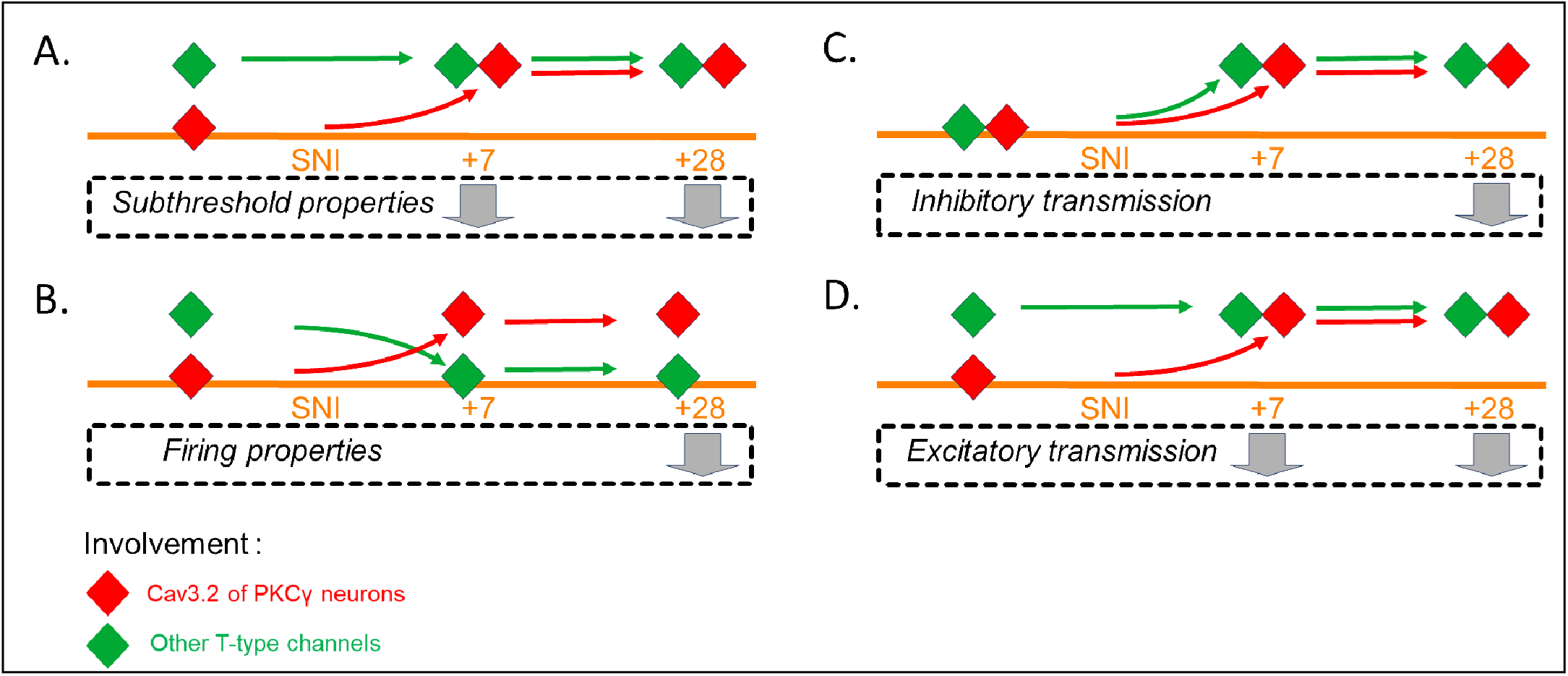
Summary of the contributions of T-type calcium channels of PKCγ neurons during mechanical allodynia. (**A-B**) The intrinsic subthreshold (**A**) and firing (**B**) properties of PKCγ neurons are diminished during early (+7: 7 days) and late (+28: 28 days) SNI as shown by the grey arrows. The contribution of Cav3.2 channels (red diamonds, inferred from comparisons between WT and KO mice) expands after SNI, unlike the contributions of the other T-type channels (green diamonds, inferred from Z944 data). (**C-D**) Remodelling of the inhibitory (**C**) and excitatory (**D**) neurotransmissions of PKCγ neurons early (+7) and late after SNI (+28). The net effect of the injury (grey arrows) was superimposed with the contributions of Cav3.2 in PKCγ neurons (red diamonds, inferred from comparisons between WT and KO mice) and of other, more distant, T-type channels (green diamonds, inferred from Z944 data).

Mechanical allodynia is characterized by a reduction in inhibitory neurotransmission, and a decrease of the synaptic inhibition on PKCγ neurons (Bardoni et al., 2013; Lu et al., 2013; Peirs et al., 2021; Petitjean et al., 2015; Qiu et al., 2022; Q. Wang et al., 2020). This disinhibition of PKCγ neurons is well-described, principally in the context of the activity evoked by primary afferent stimulation (Lu et al., 2013; Peirs et al., 2021; Q. Wang et al., 2020). Multiple mechanisms involving glutamatergic neurotransmission can also participate in central sensitization and pain processing (Inquimbert et al., 2012, 2018; Sandkühler, 2009). For instance, projection neurons of the superficial lamina I are modulated by primary afferent fibers in a T-type dependent manner (Drdla & Sandkühler, 2008; Ikeda et al., 2003) that is impaired in constitutive Cav3.2 knockout mice (Jacus et al., 2012). The glutamatergic neurotransmission of PKCγ neurons, mainly studied upon primary afferent stimulation, in not known at the synaptic level during SNI (Lu et al., 2013; Peirs et al., 2021; Q. Wang et al., 2020). Analysis of miniature currents described in the present study the status of the synapses of the PKCγ neurons during SNI. The amplitudes of mEPSC of PKCγ neurons were lowered at the early and late time points after SNI. For a comparison, previous findings described no change after spinal nerve ligation of SNI (Inquimbert et al., 2012; Zhou et al., 2011), increase during inflammation or SNI (Nelson et al., 2022; Takazawa et al., 2017), increase or decrease after sciatic nerve axotomy or sciatic chronic constriction depending on the neuronal subtype (Balasubramanyan et al., 2006; Chen et al., 2009). This decrease in mEPSC was expected to moderate the monosynaptic excitatory signaling from primary afferent neurons (Bardoni et al., 2013; Lu et al., 2013; Q. Wang et al., 2020), and might arise as a compensatory mechanism for the increase in the afferent activity during mechanical allodynia. These changes were not seen in Cav3.2-ablated PKCγ neurons, in agreement with previous findings of Cav3.2 in glutamatergic synapses in various neuronal populations (François et al., 2015; García-Caballero et al., 2014; Isope et al., 2012; Jacus et al., 2012). The evolution of mIPSC was different. The amplitudes of mIPSC were increased 7 days after SNI, while mIPSC tend to diminish in other neurons of the spinal cord (Inquimbert et al., 2018; Takazawa et al., 2017; Zhou et al., 2011) or are not changed by the pathology (Imlach et al., 2016; Zhou et al., 2011). This mechanism might strengthen the polysynaptic inhibitory signaling from primary afferent neurons (Bardoni et al., 2013; Lu et al., 2013; Q. Wang et al., 2020), and might be a compensatory mechanism for a reduction in activity of the inhibitory neurons. This increase was not seen in KO mice. Miniature IPSC appeared unchanged 28 days after SNI, and Cav3.2 was involved in this synaptic activity since its genetic ablation diminished the intervals of the mIPSC, suggesting that Cav3.2 activity was slowing the inhibitory synapse during the chronic stage of the pathology (**Fig. 9C**). These data are in agreement with roles of Cav3.2 at inhibitory synapses in various neurons including Lamina II interneurons (Candelas et al., 2019; Leresche & Lambert, 2017). In the present study, recordings of spontaneous synaptic currents provided complementary information (**Fig. 9C,D**). Indeed, spontaneous currents are a consequence of the network activity, which in the spinal cord include non-linear mechanisms such as presynaptic inhibition and excitation, shunting effects, and reverberating activity (Hachisuka et al., 2018; Ribeiro-Da-Silva & Coimbra, 1982; Zimmerman et al., 2019). The modification of IPSC was delayed, detectable as a decrease of the amplitude, 28 days after SNI only. The reduction of the amplitudes of the EPSC was observed early and late after SNI. Ablation of Cav3.2 in PKCγ neurons abolished these changes, suggesting that Cav3.2 was playing a critical role in filtering incoming synaptic inputs. It might be speculated that Cav3.2 ablation in PKCγ neurons might have similar effects on synaptic activity triggered by stimulations of some primary afferent fibers (Abraira et al., 2017). The analysis of the effects of Z944 on the spontaneous synaptic currents gave insights into the involvement of other T-type calcium channels in the synaptic networks. The T-type blocker inhibited the EPSC in PKCγ neurons of mice Sham, revealing a stimulatory role for a T-type channel at distance of the glutamatergic synapse of the PKCγ neurons. The blocker inhibited IPSC early after SNI, and no longer modulated IPSC at the later phase of the pathology, revealing a stimulatory role for a T-type channel at distance of the inhibitory synapse of the PKCγ neurons (by comparison with the effects seen in WT and KO mice). The location of these T-type channels is only hypothetical at this stage, and it is intriguing that their effects were always opposite to those of Cav3.2 of PKCγ neurons.

The net effect of Cav3.2 inhibition in PKCγ neurons on the neuronal activity or on the mice behavior could not be predicted from the present data. It was not possible to ascribe a role, either moderating or aggravating, to each of the mechanisms described here. Our data established the roles of Cav3.2 at the core of the gate control in neuropathic pain. It is striking that remodeling induced by SNI were absent in KO mice, supporting the hypothesis that Cav3.2 of PKCγ neurons might be a potential therapeutic target. Indeed, the genetic inactivation of Cav3.2 in PKCγ neurons had an antalgic effect during the early stage of the neuropathic pain, and did not change the chronic phase of the pathology. For a comparison, Z944 exerts sustained analgesic effects against chronic pain (Harding et al., 2021; LeBlanc et al., 2016) but the targets of the T-type blocker are more numerous.

Altogether, the current results indicated that a single ionic channel, Cav3.2 was involved at multiple sites of the gate control of neuropathic pain. Its restricted genetic ablation could elicit a transient antalgic effect in a mouse model of chronic pain. Next steps in the molecular analysis of this therapeutic target will be to examine its role under selective primary afferent stimulation, and recording of its putative output from the dorsal horn of the spinal card, namely layer I projection neurons. Since Cav3.2 is expressed in PKCγ neurons in humans as well as in mice, this genetic analysis will provide pre-clinical investigations required in validating this target for future therapeutical applications.

## Author contribution statement

CC, RBA, CR, MPM, PFM did the experiments; CC, EB, PFM analyzed the experiments; PF, MM provided analysis tools for patch-clamp experiments; MPM provided analysis tools for imaging; CC, JC, AF, ED, EL, EB, PFM wrote the article; EL, EB, PFM directed the work.

## ACKNOWLEDGEMENTS

We thank the staff of the RAM-iExplore and of RAM-PCEA animal facilities for assistance with transgenic mouse lines; the staff of the Montpellier Biocampus imaging core for assistance with confocal microscopy. We thank Dr E Valjent, Dr P Carroll (INM, Montpellier), Dr P Inquimbert (INCI, Strasbourg), and Dr M Antri (Neuro-Dol, Clermont-Ferrand) for helpful discussions. We thank Harald Hentschke for sharing a user-friendly interface opening abf file (https://www.mathworks.com/matlabcentral/fileexchange/6190-abfload). These studies were supported by operating grants to E Bourinet from the Agence Nationale de la Recherche (ANR LABEX ICST, PainT ANR-15-CE16-0012-01-1 : Interac-T, ANR-21-CE44-0006), Institut National de la Santé et de la Recherche Médicale, Centre National de la Recherche Scientifique, Université de Montpellier, LABEX ICST, Fondation pour la Recherche sur le cerveau (Neurodon) and la Fondation pour la Recherche Médicale (Team FRM 2015). C. Cuculière was supported by a fellowship from the LABEX ICST.

## References

Abraira, V. E., Kuehn, E. D., Chirila, A. M., Springel, M. W., Toliver, A. A., Zimmerman, A. L., Orefice, L. L., Boyle, K. A., Bai, L., Song, B. J., Bashista, K. A., O’Neill, T. G., Zhuo, J., Tsan, C., Hoynoski, J., Rutlin, M., Kus, L., Niederkofler, V., Watanabe, M., … Ginty, D. D. (2017). The Cellular and Synaptic Architecture of the Mechanosensory Dorsal Horn. Cell, 168(1-2), 295–310.e19. 10.1016/j.cell.2016.12.010

Allken, V., Chepkoech, J.-L., Einevoll, G. T., & Halnes, G. (2014). The Subcellular Distribution of T-Type Ca2+ Channels in Interneurons of the Lateral Geniculate Nucleus. PLOS ONE, 9(9), e107780. 10.1371/journal.pone.0107780

Baccam, N., Alonso, G., Costecalde, T., Fontanaud, P., Molino, F., Robinson, I. C. A. F., Mollard, P., & Méry, P.-F. (2007). Dual-Level Afferent Control of Growth Hormone-Releasing Hormone (GHRH) Neurons in GHRH–Green Fluorescent Protein Transgenic Mice. Journal of Neuroscience, 27(7), 1631–1641. 10.1523/JNEUROSCI.2693-06.2007

Balasubramanyan, S., Stemkowski, P. L., Stebbing, M. J., & Smith, P. A. (2006). Sciatic chronic constriction injury produces cell-type-specific changes in the electrophysiological properties of rat substantia gelatinosa neurons. Journal of Neurophysiology, 96(2), 579–590. 10.1152/jn.00087.2006

Bardoni, R., Takazawa, T., Tong, C.-K., Choudhury, P., Scherrer, G., & MacDermott, A. B. (2013). Pre-and postsynaptic inhibitory control in the spinal cord dorsal horn. Annals of the New York Academy of Sciences, 1279(1), 90–96. 10.1111/nyas.12056

Bosch, M. A., Hou, J., Fang, Y., Kelly, M. J., & RØnnekleiv, O. K. (2009). 17β-estradiol regulation of the mRNA expression of t-type calcium channel subunits : Role of estrogen receptor α and estrogen receptor β. Journal of Comparative Neurology, 512(3), 347–358. 10.1002/cne.21901

Bourinet, E., Alloui, A., Monteil, A., Barrère, C., Couette, B., Poirot, O., Pages, A., McRory, J., Snutch, T. P., Eschalier, A., & Nargeot, J. (2005). Silencing of the Cav3.2 T-type calcium channel gene in sensory neurons demonstrates its major role in nociception. The EMBO Journal, 24(2), Article 2. 10.1038/sj.emboj.7600515

Bourinet, E., Francois, A., & Laffray, S. (2016). T-type calcium channels in neuropathic pain. PAIN, 157, S15. 10.1097/j.pain.0000000000000469

Boyle, K. A., Gradwell, M. A., Yasaka, T., Dickie, A. C., Polgár, E., Ganley, R. P., Orr, D. P. H., Watanabe, M., Abraira, V. E., Kuehn, E. D., Zimmerman, A. L., Ginty, D. D., Callister, R. J., Graham, B. A., & Hughes, D. I. (2019). Defining a Spinal Microcircuit that Gates Myelinated Afferent Input : Implications for Tactile Allodynia. Cell Reports, 28(2), 526–540.e6. 10.1016/j.celrep.2019.06.040

Cain, S. M., & Snutch, T. P. (2010). Contributions of T-type calcium channel isoforms to neuronal firing. Channels (Austin, Tex.), 4(6), Article 6. 10.4161/chan.4.6.14106

Candelas, M., Reynders, A., Arango-Lievano, M., Neumayer, C., Fruquière, A., Demes, E., Hamid, J., Lemmers, C., Bernat, C., Monteil, A., Compan, V., Laffray, S., Inquimbert, P., Le Feuvre, Y., Zamponi, G. W., Moqrich, A., Bourinet, E., & Méry, P.-F. (2019). Cav3.2 T-type calcium channels shape electrical firing in mouse Lamina II neurons. Scientific Reports, 9(1), Article 1. 10.1038/s41598-019-39703-3

Chaplan, S. R., Bach, F. W., Pogrel, J. W., Chung, J. M., & Yaksh, T. L. (1994). Quantitative assessment of tactile allodynia in the rat paw. Journal of Neuroscience Methods, 53(1), 55–63. 10.1016/0165-0270(94)90144-9

Chen, S.-R., Cai, Y.-Q., & Pan, H.-L. (2009). Plasticity and emerging role of BKCa channels in nociceptive control in neuropathic pain. Journal of Neurochemistry, 110(1), 352–362. 10.1111/j.1471-4159.2009.06138.x

Clements, J. D., & Bekkers, J. M. (1997). Detection of spontaneous synaptic events with an optimally scaled template. Biophysical Journal, 73(1), 220–229. 10.1016/S0006-3495(97)78062-7

Decosterd, I., & Woolf, C. J. (2000). Spared nerve injury : An animal model of persistent peripheral neuropathic pain. Pain, 87(2), 149–158. 10.1016/S0304-3959(00)00276-1

Deleuze, C., David, F., Béhuret, S., Sadoc, G., Shin, H.-S., Uebele, V. N., Renger, J. J., Lambert, R. C., Leresche, N., & Bal, T. (2012). T-Type Calcium Channels Consolidate Tonic Action Potential Output of Thalamic Neurons to Neocortex. Journal of Neuroscience, 32(35), 12228–12236. 10.1523/JNEUROSCI.1362-12.2012

Dou, Y., Xia, J., Gao, R., Gao, X., Munoz, F. M., Wei, D., Tian, Y., Barrett, J. E., Ajit, S., Meucci, O., Putney, J. W., Dai, Y., & Hu, H. (2018). Orai1 Plays a Crucial Role in Central Sensitization by Modulating Neuronal Excitability. Journal of Neuroscience, 38(4), 887–900. 10.1523/JNEUROSCI.3007-17.2017

Drdla, R., & Sandkühler, J. (2008). Long-Term Potentiation at C-Fibre Synapses by Low-Level Presynaptic Activity in vivo. Molecular Pain, 4, 1744–8069-4-18. 10.1186/1744-8069-4-18

Dreyfus, F. M., Tscherter, A., Errington, A. C., Renger, J. J., Shin, H.-S., Uebele, V. N., Crunelli, V., Lambert, R. C., & Leresche, N. (2010). Selective T-Type Calcium Channel Block in Thalamic Neurons Reveals Channel Redundancy and Physiological Impact of ITwindow. Journal of Neuroscience, 30(1), 99–109. 10.1523/JNEUROSCI.4305-09.2010

Dumenieu, M., Senkov, O., Mironov, A., Bourinet, E., Kreutz, M. R., Dityatev, A., Heine, M., Bikbaev, A., & Lopez-Rojas, J. (2018). The Low-Threshold Calcium Channel Cav3.2 Mediates Burst Firing of Mature Dentate Granule Cells. Cerebral Cortex, 28(7), 2594–2609. 10.1093/cercor/bhy084

Eickhoff, A., Tjaden, J., Stahlke, S., Vorgerd, M., Theis, V., Matschke, V., & Theiss, C. (2022). Effects of progesterone on T-type-Ca2+-channel expression in Purkinje cells. Neural Regeneration Research, 17(11), 2465–2471. 10.4103/1673-5374.339008

Engbers, J. D. T., Anderson, D., Asmara, H., Rehak, R., Mehaffey, W. H., Hameed, S., McKay, B. E., Kruskic, M., Zamponi, G. W., & Turner, R. W. (2012). Intermediate conductance calcium-activated potassium channels modulate summation of parallel fiber input in cerebellar Purkinje cells. Proceedings of the National Academy of Sciences, 109(7), 2601–2606. 10.1073/pnas.1115024109

Engbers, J. D. T., Anderson, D., Tadayonnejad, R., Mehaffey, W. H., Molineux, M. L., & Turner, R. W. (2011). Distinct roles for IT and IH in controlling the frequency and timing of rebound spike responses. The Journal of Physiology, 589(22), 5391–5413. 10.1113/jphysiol.2011.215632

Fayad, S. L., Ourties, G., Le Gac, B., Jouffre, B., Lamoine, S., Fruquière, A., Laffray, S., Gasmi, L., Cauli, B., Mallet, C., Bourinet, E., Bessaih, T., Lambert, R. C., & Leresche, N. (2022). Centrally expressed Cav3.2 T-type calcium channel is critical for the initiation and maintenance of neuropathic pain. eLife, 11, e79018. 10.7554/eLife.79018

François, A., Schüetter, N., Laffray, S., Sanguesa, J., Pizzoccaro, A., Dubel, S., Mantilleri, A., Nargeot, J., Noël, J., Wood, J. N., Moqrich, A., Pongs, O., & Bourinet, E. (2015). The Low-Threshold Calcium Channel Cav3.2 Determines Low-Threshold Mechanoreceptor Function. Cell Reports, 10(3), Article 3. 10.1016/j.celrep.2014.12.042

Garcia, M. M., Goicoechea, C., Avellanal, M., Traseira, S., Martín, M. I., & Sánchez-Robles, E. M. (2019). Comparison of the antinociceptive profiles of morphine and oxycodone in two models of inflammatory and osteoarthritic pain in rat. European Journal of Pharmacology, 854, 109–118. 10.1016/j.ejphar.2019.04.011

García-Caballero, A., Gadotti, V. M., Stemkowski, P., Weiss, N., Souza, I. A., Hodgkinson, V., Bladen, C., Chen, L., Hamid, J., Pizzoccaro, A., Deage, M., François, A., Bourinet, E., & Zamponi, G. W. (2014). The Deubiquitinating Enzyme USP5 Modulates Neuropathic and Inflammatory Pain by Enhancing Cav3.2 Channel Activity. Neuron, 83(5), 1144–1158. 10.1016/j.neuron.2014.07.036

Gassner, M., Leitner, J., Gruber-Schoffnegger, D., Forsthuber, L., & Sandkühler, J. (2013). Properties of spinal lamina III GABAergic neurons in naïve and in neuropathic mice. *European Journal of Pain (London*, England), 17(8), 1168–1179. 10.1002/j.1532-2149.2013.00294.x

Ghazisaeidi, S., Muley, M. M., & Salter, M. W. (2023). Neuropathic Pain : Mechanisms, Sex Differences, and Potential Therapies for a Global Problem. Annual Review of Pharmacology and Toxicology, 63, 565–583. 10.1146/annurev-pharmtox-051421-112259

Grudt, T. J., & Perl, E. R. (2002). Correlations between neuronal morphology and electrophysiological features in the rodent superficial dorsal horn. The Journal of Physiology, 540(1), 189–207. 10.1113/jphysiol.2001.012890

Hachisuka, J., Omori, Y., Chiang, M. C., Gold, M. S., Koerber, H. R., & Ross, S. E. (2018). Wind-up in lamina I spinoparabrachial neurons : A role for reverberatory circuits. Pain, 159(8), 1484–1493. 10.1097/j.pain.0000000000001229

Harding, E. K., Dedek, A., Bonin, R. P., Salter, M. W., Snutch, T. P., & Hildebrand, M. E. (2021). The T-type calcium channel antagonist, Z944, reduces spinal excitability and pain hypersensitivity. British Journal of Pharmacology, 178(17), 3517–3532. 10.1111/bph.15498

He, X., Liu, P., Zhang, X., Jiang, Z., Gu, N., Wang, Q., & Lu, Y. (2021). Molecular and Electrophysiological Characterization of Dorsal Horn Neurons in a GlyT2-iCre-tdTomato Mouse Line. Journal of Pain Research, 14, 907–921. 10.2147/JPR.S296940

Hu, B. (1995). Cellular basis of temporal synaptic signalling : An in vitro electrophysiological study in rat auditory thalamus. The Journal of Physiology, 483(1), 167–182. 10.1113/jphysiol.1995.sp020576

Ikeda, H., Heinke, B., Ruscheweyh, R., & Sandkühler, J. (2003). Synaptic Plasticity in Spinal Lamina I Projection Neurons That Mediate Hyperalgesia. Science, 299(5610), 1237–1240. 10.1126/science.1080659

Imlach, W. L., Bhola, R. F., Mohammadi, S. A., & Christie, M. J. (2016). Glycinergic dysfunction in a subpopulation of dorsal horn interneurons in a rat model of neuropathic pain. Scientific Reports, 6(1), Article 1. 10.1038/srep37104

Inquimbert, P., Bartels, K., Babaniyi, O. B., Barrett, L. B., Tegeder, I., & Scholz, J. (2012). Peripheral nerve injury produces a sustained shift in the balance between glutamate release and uptake in the dorsal horn of the spinal cord. PAIN®, 153(12), 2422–2431. 10.1016/j.pain.2012.08.011

Inquimbert, P., Moll, M., Latremoliere, A., Tong, C.-K., Whang, J., Sheehan, G. F., Smith, B. M., Korb, E., Athié, M. C. P., Babaniyi, O., Ghasemlou, N., Yanagawa, Y., Allis, C. D., Hof, P. R., & Scholz, J. (2018). NMDA Receptor Activation Underlies the Loss of Spinal Dorsal Horn Neurons and the Transition to Persistent Pain after Peripheral Nerve Injury. Cell Reports, 23(9), 2678–2689. 10.1016/j.celrep.2018.04.107

Inyang, K., Szabo-Pardi, T., & Price, T. (2016). (309) Treatment of Chronic pain : Long term effects of Metformin on chronic neuropathic pain and microglial activation. The Journal of Pain, 17(4), S53. 10.1016/j.jpain.2016.01.216

Ippolito, D. L., Xu, M., Bruchas, M. R., Wickman, K., & Chavkin, C. (2005). Tyrosine Phosphorylation of Kir3.1 in Spinal Cord Is Induced by Acute Inflammation, Chronic Neuropathic Pain, and Behavioral Stress *. Journal of Biological Chemistry, 280(50), 41683–41693. 10.1074/jbc.M507069200

Isope, P., Hildebrand, M. E., & Snutch, T. P. (2012). Contributions of T-Type Voltage-Gated Calcium Channels to Postsynaptic Calcium Signaling within Purkinje Neurons. The Cerebellum, 11(3), 651–665. 10.1007/s12311-010-0195-4

Jacus, M. O., Uebele, V. N., Renger, J. J., & Todorovic, S. M. (2012). Presynaptic CaV3.2 channels regulate excitatory neurotransmission in nociceptive dorsal horn neurons. The Journal of Neuroscience, 32(27), Article 27. 10.1523/JNEUROSCI.0068-12.2012

Jaggi, A. S., Jain, V., & Singh, N. (2011). Animal models of neuropathic pain. Fundamental & Clinical Pharmacology, 25(1), 1–28. 10.1111/j.1472-8206.2009.00801.x

Jeon, S.-M., Chang, D., Geske, A., Ginty, D. D., & Caterina, M. J. (2021). Sex-Dependent Reduction in Mechanical Allodynia in the Sural-Sparing Nerve Injury Model in Mice Lacking Merkel Cells. Journal of Neuroscience, 41(26), 5595–5619. 10.1523/JNEUROSCI.1668-20.2021

Kerckhove, N., Mallet, C., François, A., Boudes, M., Chemin, J., Voets, T., Bourinet, E., Alloui, A., & Eschalier, A. (2014). Cav3.2 calcium channels : The key protagonist in the supraspinal effect of paracetamol. PAIN®, 155(4), 764–772. 10.1016/j.pain.2014.01.015

Kopach, O., Krotov, V., Belan, P., & Voitenko, N. (2015). Inflammatory-induced changes in synaptic drive and postsynaptic AMPARs in lamina II dorsal horn neurons are cell-type specific. PAIN, 156(3), 428. 10.1097/01.j.pain.0000460318.65734.00

Kovács, K., Sík, A., Ricketts, C., & Timofeev, I. (2010). Subcellular distribution of low-voltage activated T-type Ca2+ channel subunits (Cav3.1 and Cav3.3) in reticular thalamic neurons of the cat. Journal of Neuroscience Research, 88(2), 448–460. 10.1002/jnr.22200

Ku, W., & Schneider, S. P. (2011). Multiple T-type Ca2+ current subtypes in electrophysiologically characterized hamster dorsal horn neurons : Possible role in spinal sensory integration. Journal of Neurophysiology, 106(5), 2486–2498. 10.1152/jn.01083.2010

Kuhn, J. A., Vainchtein, I. D., Braz, J., Hamel, K., Bernstein, M., Craik, V., Dahlgren, M. W., Ortiz-Carpena, J., Molofsky, A. B., Molofsky, A. V., & Basbaum, A. I. (2021). Regulatory T-cells inhibit microglia-induced pain hypersensitivity in female mice. eLife, 10, e69056. 10.7554/eLife.69056

Latham, J. R., Pathirathna, S., Jagodic, M. M., Joo Choe, W., Levin, M. E., Nelson, M. T., Yong Lee, W., Krishnan, K., Covey, D. F., Todorovic, S. M., & Jevtovic-Todorovic, V. (2009). Selective T-Type Calcium Channel Blockade Alleviates Hyperalgesia in ob/ob Mice. Diabetes, 58(11), 2656–2665. 10.2337/db08-1763

LeBlanc, B. W., Lii, T. R., Huang, J. J., Chao, Y.-C., Bowary, P. M., Cross, B. S., Lee, M. S., Vera-Portocarrero, L. P., & Saab, C. Y. (2016). T-type calcium channel blocker Z944 restores cortical synchrony and thalamocortical connectivity in a rat model of neuropathic pain. PAIN, 157(1), 255. 10.1097/j.pain.0000000000000362

Lee, J.-H., Gomora, J. C., Cribbs, L. L., & Perez-Reyes, E. (1999). Nickel Block of Three Cloned T-Type Calcium Channels : Low Concentrations Selectively Block α1H. Biophysical Journal, 77(6), 3034–3042. 10.1016/S0006-3495(99)77134-1

Leresche, N., & Lambert, R. C. (2017). T-type calcium channels in synaptic plasticity. Channels, 11(2), 121–139. 10.1080/19336950.2016.1238992

Liao, Y.-F., Tsai, M.-L., Chen, C.-C., & Yen, C.-T. (2011). Involvement of the Cav3.2 T-Type Calcium Channel in Thalamic Neuron Discharge Patterns. Molecular Pain, 7, 1744–8069-7-43. 10.1186/1744-8069-7-43

Lipkin, A. M., Cunniff, M. M., Spratt, P. W. E., Lemke, S. M., & Bender, K. J. (2021). Functional Microstructure of CaV-Mediated Calcium Signaling in the Axon Initial Segment. Journal of Neuroscience, 41(17), 3764–3776. 10.1523/JNEUROSCI.2843-20.2021

Llinás, R. R., & Steriade, M. (2006). Bursting of Thalamic Neurons and States of Vigilance. Journal of Neurophysiology, 95(6), 3297–3308. 10.1152/jn.00166.2006

Lu, Y., Dong, H., Gao, Y., Gong, Y., Ren, Y., Gu, N., Zhou, S., Xia, N., Sun, Y.-Y., Ji, R.-R., & Xiong, L. (2013). A feed-forward spinal cord glycinergic neural circuit gates mechanical allodynia. The Journal of Clinical Investigation, 123(9), 4050–4062. 10.1172/JCI70026

Ma, X., Miraucourt, L. S., Qiu, H., Sharif-Naeini, R., & Khadra, A. (2023). Modulation of SK Channels via Calcium Buffering Tunes Intrinsic Excitability of Parvalbumin Interneurons in Neuropathic Pain : A Computational and Experimental Investigation. Journal of Neuroscience, 43(31), 5608–5622. 10.1523/JNEUROSCI.0426-23.2023

Madisen, L., Zwingman, T. A., Sunkin, S. M., Oh, S. W., Zariwala, H. A., Gu, H., Ng, L. L., Palmiter, R. D., Hawrylycz, M. J., Jones, A. R., Lein, E. S., & Zeng, H. (2010). A robust and high-throughput Cre reporting and characterization system for the whole mouse brain. Nature Neuroscience, 13(1), Article 1. 10.1038/nn.2467

Marger, F., Gelot, A., Alloui, A., Matricon, J., Ferrer, J. F. S., Barrere, C., Pizzoccaro, A., Muller, E., Nargeot, J., Snutch, T. P., Eschalier, A., Bourinet, E., & Ardid, D. (2011). T-type calcium channels contribute to colonic hypersensitivity in a rat model of irritable bowel syndrome. Proceedings of the National Academy of Sciences, 108(27), Article 27. 10.1073/pnas.1100869108

Melnick, I. V., Santos, S. F. A., & Safronov, B. V. (2004). Mechanism of spike frequency adaptation in substantia gelatinosa neurones of rat. The Journal of Physiology, 559(Pt 2), 383–395. 10.1113/jphysiol.2004.066415

Messinger, R. B., Naik, A. K., Jagodic, M. M., Nelson, M. T., Lee, W. Y., Choe, W. J., Orestes, P., Latham, J. R., Todorovic, S. M., & Jevtovic-Todorovic, V. (2009). In vivo silencing of the CaV3.2 T-type calcium channels in sensory neurons alleviates hyperalgesia in rats with streptozocin-induced diabetic neuropathy. PAIN®, 145(1), 184–195. 10.1016/j.pain.2009.06.012

Mogil, J. S. (2020). Qualitative sex differences in pain processing : Emerging evidence of a biased literature. Nature Reviews Neuroscience, 21(7), Article 7. 10.1038/s41583-020-0310-6

Molineux, M. L., McRory, J. E., McKay, B. E., Hamid, J., Mehaffey, W. H., Rehak, R., Snutch, T. P., Zamponi, G. W., & Turner, R. W. (2006). Specific T-type calcium channel isoforms are associated with distinct burst phenotypes in deep cerebellar nuclear neurons. Proceedings of the National Academy of Sciences, 103(14), 5555–5560. 10.1073/pnas.0601261103

Nakagawa, T., Yasaka, T., Nakashima, N., Takeya, M., Oshita, K., Tsuda, M., Yamaura, K., & Takano, M. (2020). Expression of the pacemaker channel HCN4 in excitatory interneurons in the dorsal horn of the murine spinal cord. Molecular Brain, 13(1), 127. 10.1186/s13041-020-00666-6

Nam, G. (2018). T-type calcium channel blockers : A patent review (2012-2018). Expert Opinion on Therapeutic Patents, 28(12), 883–901. 10.1080/13543776.2018.1541982

Nelson, T. S., Sinha, G. P., Santos, D. F. S., Jukkola, P., Prasoon, P., Winter, M. K., McCarson, K. E., Smith, B. N., & Taylor, B. K. (2022). Spinal neuropeptide Y Y1 receptor-expressing neurons are a pharmacotherapeutic target for the alleviation of neuropathic pain. Proceedings of the National Academy of Sciences, 119(46), e2204515119. 10.1073/pnas.2204515119

Neumann, S., Braz, J. M., Skinner, K., Llewellyn-Smith, I. J., & Basbaum, A. I. (2008). Innocuous, Not Noxious, Input Activates PKCγ Interneurons of the Spinal Dorsal Horn via Myelinated Afferent Fibers. The Journal of Neuroscience, 28(32), 7936–7944. 10.1523/JNEUROSCI.1259-08.2008

Paige, C., Plasencia-Fernandez, I., Kume, M., Papalampropoulou-Tsiridou, M., Lorenzo, L.-E., David, E. T., He, L., Mejia, G. L., Driskill, C., Ferrini, F., Feldhaus, A. L., Garcia-Martinez, L. F., Akopian, A. N., Koninck, Y. D., Dussor, G., & Price, T. J. (2022). A Female-Specific Role for Calcitonin Gene-Related Peptide (CGRP) in Rodent Pain Models. Journal of Neuroscience, 42(10), 1930–1944. 10.1523/JNEUROSCI.1137-21.2022

Pathirathna, S., Brimelow, B. C., Jagodic, M. M., Krishnan, K., Jiang, X., Zorumski, C. F., Mennerick, S., Covey, D. F., Todorovic, S. M., & Jevtovic-Todorovic, V. (2005). New evidence that both T-type calcium channels and GABAA channels are responsible for the potent peripheral analgesic effects of 5alpha-reduced neuroactive steroids. Pain, 114(3), 429–443. 10.1016/j.pain.2005.01.009

Peirs, C., Williams, S.-P. G., Zhao, X., Arokiaraj, C. M., Ferreira, D. W., Noh, M.-C., Smith, K. M., Halder, P., Corrigan, K. A., Gedeon, J. Y., Lee, S. J., Gatto, G., Chi, D., Ross, S. E., Goulding, M., & Seal, R. P. (2021). Mechanical Allodynia Circuitry in the Dorsal Horn Is Defined by the Nature of the Injury. Neuron, 109(1), 73–90.e7. 10.1016/j.neuron.2020.10.027

Peirs, C., Williams, S.-P. G., Zhao, X., Walsh, C. E., Gedeon, J. Y., Cagle, N. E., Goldring, A. C., Hioki, H., Liu, Z., Marell, P. S., & Seal, R. P. (2015). Dorsal Horn Circuits for Persistent Mechanical Pain. Neuron, 87(4), 797–812. 10.1016/j.neuron.2015.07.029

Petitjean, H., Pawlowski, S. A., Fraine, S. L., Sharif, B., Hamad, D., Fatima, T., Berg, J., Brown, C. M., Jan, L.-Y., Ribeiro-da-Silva, A., Braz, J. M., Basbaum, A. I., & Sharif-Naeini, R. (2015). Dorsal Horn Parvalbumin Neurons Are Gate-Keepers of Touch-Evoked Pain after Nerve Injury. Cell reports, 13(6), Article 6. 10.1016/j.celrep.2015.09.080

Prescott, S. A., & Koninck, Y. D. (2002). Four cell types with distinctive membrane properties and morphologies in lamina I of the spinal dorsal horn of the adult rat. The Journal of Physiology, 539(3), 817–836. 10.1113/jphysiol.2001.013437

Qiu, H., Miraucourt, L., Petitjean, H., Davidova, A., Levesque-Damphousse, P., Estall, J. L., & Sharif-Naeini, R. (2022). *Parvalbumin protein controls inhibitory tone in the spinal cord* [Preprint]. Neuroscience. 10.1101/2022.09.15.508019

Ratté, S., Lankarany, M., Rho, Y.-A., Patterson, A., & Prescott, S. A. (2015). Subthreshold membrane currents confer distinct tuning properties that enable neurons to encode the integral or derivative of their input. Frontiers in Cellular Neuroscience, 8. https://www.frontiersin.org/articles/10.3389/fncel.2014.00452

Ribeiro-Da-Silva, A., & Coimbra, A. (1982). Two types of synaptic glomeruli and their distribution in laminae I-III of the rat spinal cord. The Journal of Comparative Neurology, 209(2), 176–186. 10.1002/cne.902090205

Ruscheweyh, R., & Sandkühler, J. (2002). Role of kainate receptors in nociception. Brain Research Reviews, 40(1), 215–222. 10.1016/S0165-0173(02)00203-5

Russ, D. E., Cross, R. B. P., Li, L., Koch, S. C., Matson, K. J. E., Yadav, A., Alkaslasi, M. R., Lee, D. I., Le Pichon, C. E., Menon, V., & Levine, A. J. (2021). A harmonized atlas of mouse spinal cord cell types and their spatial organization. Nature Communications, 12(1), 5722. 10.1038/s41467-021-25125-1

Sandkühler, J. (2009). Models and mechanisms of hyperalgesia and allodynia. Physiological Reviews, 89(2), 707–758. 10.1152/physrev.00025.2008

Schindelin, J., Arganda-Carreras, I., Frise, E., Kaynig, V., Longair, M., Pietzsch, T., Preibisch, S., Rueden, C., Saalfeld, S., Schmid, B., Tinevez, J.-Y., White, D. J., Hartenstein, V., Eliceiri, K., Tomancak, P., & Cardona, A. (2012). Fiji—An Open Source platform for biological image analysis. Nature methods, 9(7), Article 7. 10.1038/nmeth.2019

Schoffnegger, D., Heinke, B., Sommer, C., & Sandkühler, J. (2006). Physiological properties of spinal lamina II GABAergic neurons in mice following peripheral nerve injury. The Journal of Physiology, 577(Pt 3), 869–878. 10.1113/jphysiol.2006.118034

Shen, F.-Y., Chen, Z.-Y., Zhong, W., Ma, L.-Q., Chen, C., Yang, Z.-J., Xie, W.-L., & Wang, Y.-W. (2015). Alleviation of Neuropathic Pain by Regulating T-Type Calcium Channels in Rat Anterior Cingulate Cortex. Molecular Pain, 11, s12990–015-0008. 10.1186/s12990-015-0008-3

Sinha, G. P., Prasoon, P., Smith, B. N., & Taylor, B. K. (2021). Fast A-type currents shape a rapidly adapting form of delayed short latency firing of excitatory superficial dorsal horn neurons that express the neuropeptide Y Y1 receptor. The Journal of Physiology, 599(10), 2723–2750. 10.1113/JP281033

Smith, K. M., Boyle, K. A., Madden, J. F., Dickinson, S. A., Jobling, P., Callister, R. J., Hughes, D. I., & Graham, B. A. (2015). Functional heterogeneity of calretinin-expressing neurons in the mouse superficial dorsal horn : Implications for spinal pain processing. The Journal of Physiology, 593(19), 4319–4339. 10.1113/JP270855

Snutch, T. P., & Zamponi, G. W. (2018). Recent advances in the development of T-type calcium channel blockers for pain intervention. British Journal of Pharmacology, 175(12), 2375–2383. 10.1111/bph.13906

Sun, H., Xu, J., Della Penna, K. B., Benz, R. J., Kinose, F., Holder, D. J., Koblan, K. S., Gerhold, D. L., & Wang, H. (2002). Dorsal horn-enriched genes identified by DNA microarray, in situ hybridization and immunohistochemistry. BMC Neuroscience, 3(1), 11. 10.1186/1471-2202-3-11

Tadros, M. A., Harris, B. M., Anderson, W. B., Brichta, A. M., Graham, B. A., & Callister, R. J. (2012). Are all spinal segments equal : Intrinsic membrane properties of superficial dorsal horn neurons in the developing and mature mouse spinal cord. The Journal of Physiology, 590(10), 2409–2425. 10.1113/jphysiol.2012.227389

Takazawa, T., Choudhury, P., Tong, C.-K., Conway, C. M., Scherrer, G., Flood, P. D., Mukai, J., & MacDermott, A. B. (2017). Inhibition Mediated by Glycinergic and GABAergic Receptors on Excitatory Neurons in Mouse Superficial Dorsal Horn Is Location-Specific but Modified by Inflammation. The Journal of Neuroscience: The Official Journal of the Society for Neuroscience, 37(9), 2336–2348. 10.1523/JNEUROSCI.2354-16.2017

Temmermand, R., Barrett, J. E., & Fontana, A. C. K. (2022). Glutamatergic systems in neuropathic pain and emerging non-opioid therapies. Pharmacological Research, 185, 106492. 10.1016/j.phrs.2022.106492

Tran, M., Braz, J. M., Hamel, K., Kuhn, J., Todd, A. J., & Basbaum, A. I. (2020). Ablation of spinal cord estrogen receptor α-expressing interneurons reduces chemically induced modalities of pain and itch. The Journal of Comparative Neurology, 528(10), 1629–1643. 10.1002/cne.24847

Tringham, E., Powell, K. L., Cain, S. M., Kuplast, K., Mezeyova, J., Weerapura, M., Eduljee, C., Jiang, X., Smith, P., Morrison, J.-L., Jones, N. C., Braine, E., Rind, G., Fee-Maki, M., Parker, D., Pajouhesh, H., Parmar, M., O’Brien, T. J., & Snutch, T. P. (2012). T-Type Calcium Channel Blockers That Attenuate Thalamic Burst Firing and Suppress Absence Seizures. Science Translational Medicine, 4(121), 121ra19–121ra19. 10.1126/scitranslmed.3003120

Wang, G., Bochorishvili, G., Chen, Y., Salvati, K. A., Zhang, P., Dubel, S. J., Perez-Reyes, E., Snutch, T. P., Stornetta, R. L., Deisseroth, K., Erisir, A., Todorovic, S. M., Luo, J.-H., Kapur, J., Beenhakker, M. P., & Zhu, J. J. (2015). CaV3.2 calcium channels control NMDA receptor-mediated transmission : A new mechanism for absence epilepsy. Genes & Development, 29(14), 1535–1551. 10.1101/gad.260869.115

Wang, Q., Zhang, X., He, X., Du, S., Jiang, Z., Liu, P., Qi, L., Liang, C., Gu, N., & Lu, Y. (2020). Synaptic Dynamics of the Feed-forward Inhibitory Circuitry Gating Mechanical Allodynia in Mice. Anesthesiology, 132(5), 1212–1228. 10.1097/ALN.0000000000003194

Wang, R., & Lewin, G. R. (2011). The Cav3.2 T-type calcium channel regulates temporal coding in mouse mechanoreceptors. The Journal of Physiology, 589(9), 2229–2243. 10.1113/jphysiol.2010.203463

Yadav, A., Matson, K. J. E., Li, L., Hua, I., Petrescu, J., Kang, K., Alkaslasi, M. R., Lee, D. I., Hasan, S., Galuta, A., Dedek, A., Ameri, S., Parnell, J., Alshardan, M. M., Qumqumji, F. A., Alhamad, S. M., Wang, A. P., Poulen, G., Lonjon, N., … Levine, A. J. (2023). A cellular taxonomy of the adult human spinal cord. Neuron, 111(3), 328–344.e7. 10.1016/j.neuron.2023.01.007

Yoo, S., Santos, C., Reynders, A., Ugolini, S., Rossignol, R., Khallouqi, A. E., Springael, J.-Y., Parmentier, M., Saurin, A. J., Goaillard, J.-M., Castets, F., Clerc, N., & Moqrich, A. (s. d.). TAFA4 relieves injury-induced mechanical hypersensitivity through LDL receptors and modulation of spinal A-type K+ current. OPEN ACCESS.

Zamponi, G. W., Striessnig, J., Koschak, A., & Dolphin, A. C. (2015). The Physiology, Pathology, and Pharmacology of Voltage-Gated Calcium Channels and Their Future Therapeutic Potential. Pharmacological Reviews, 67(4), 821–870. 10.1124/pr.114.009654

Zhang, C., Bosch, M. A., Rick, E. A., Kelly, M. J., & Rønnekleiv, O. K. (2009). 17β-Estradiol Regulation of T-Type Calcium Channels in Gonadotropin-Releasing Hormone Neurons. Journal of Neuroscience, 29(34), 10552–10562. 10.1523/JNEUROSCI.2962-09.2009

Zhi, Y.-R., Cao, F., Su, X.-J., Gao, S.-W., Zheng, H.-N., Jiang, J.-Y., Su, L., Liu, J., Wang, Y., Zhang, Y., & Zhang, Y. (2022). The T-Type Calcium Channel Cav3.2 in Somatostatin Interneurons in Spinal Dorsal Horn Participates in Mechanosensation and Mechanical Allodynia in Mice. Frontiers in Cellular Neuroscience, 16. https://www.frontiersin.org/articles/10.3389/fncel.2022.875726

Zhou, H.-Y., Chen, S.-R., Chen, H., & Pan, H.-L. (2011). Functional Plasticity of Group II Metabotropic Glutamate Receptors in Regulating Spinal Excitatory and Inhibitory Synaptic Input in Neuropathic Pain. Journal of Pharmacology and Experimental Therapeutics, 336(1), 254–264. 10.1124/jpet.110.173112

Zimmerman, A. L., Kovatsis, E. M., Pozsgai, R. Y., Tasnim, A., Zhang, Q., & Ginty, D. D. (2019). Distinct Modes of Presynaptic Inhibition of Cutaneous Afferents and Their Functions in Behavior. Neuron, 102(2), 420–434.e8. 10.1016/j.neuron.2019.02.002

