## Supplementary figures 1-5 for "T-type calcium channels participate in intrinsic and synaptic activity of PKCγ neurons of the dorsal horn of the spinal cord during chronic pain"

**Célia Cuculière[1,2,3,4], Rachel Bourdon-Alonzeau[1,2,3,4], Coline Rulhe[1,2,3,4], Jean Chemin[1,2,3,4], Amaury François[1,2,3,4], Marie-Pierre Blanchard[5], Pierre Fontanaud[1,2,3,4], Emmanuel Deval[6], Matteo Mangoni[1,2,3,4], Eric Lingueglia[6], Emmanuel Bourinet[1,2,3,4*], Pierre-François Méry[1,2,3,4 *]**

**Addresses:** [1] Laboratories of Excellence - Ion Channel Science and Therapeutics, Montpellier, France, [2] Inserm U-1191, Montpellier, France; [3] CNRS UMR 5203, Institut de Génomique Fonctionnelle, Montpellier, France; [4] Université Montpellier, Montpellier, France; [5] Montpellier Ressources Imagerie, BioCampus, Université de Montpellier, CNRS, INSERM, Montpellier, France; [6] Université Côte d'Azur, CNRS, Institut de Pharmacologie Moléculaire et Cellulaire (IPMC), LabEx Ion Channel Science and Therapeutics (ICST), FHU Innovative Solutions in Refractory Chronic Pain (InovPain), France.

**Supplementary Figure 1. Subcellular distribution and functional properties of Cav3.2 in PKCγ neurons.**

**(A)** Immunostainings of a whole lumbar spinal cord from adult PKCγ-Cre$^{ERT2}$ x Ai14-tdTomato with a homozygous Cav3.2$^{GFP\text{-}Flox}$ KI background. This representative example shows the PKCγ expressing cells (red) in a Sytox (green) counterstaining. **(B)** Immunostaining for GFP (green) and PKCγ (red), showing the subcellular localization of Cav3.2-GFP in PKCγ expressing neurons (yellow signal) in the dorsal horn of the lumbar spinal cord. The majority of GFP is found in the soma and primary dendrites, and can be seen in presumptive synaptic buttons (yellow arrowheads). It was absent of axon initial segment marked by Ankyrin G (white). Scale bars: **A** (50 µm), **B** (10 µm). **(C)** Family of typical T-type calcium currents recorded in a PKCγ neuron. Currents were elicited at increasing membrane potentials, as indicated, from a holding potential of -100 mV.

**Supplementary Figure 2. Subthreshold properties of PKCγ neurons**

**(A)** Proportions of after-hyperpolarisations responses (rebound, hyperpolarizing, flat) recorded in PKCγ neurons of WT and SNI KO mice with the whole-cell patch clamp technique (see Methods for details). **(B)** Maximal rebound amplitude recorded in current-clamp experiments in naïve and after Sham surgery in WT mice. Number of experiments were: 30 (Naïve), 30 (Sham 7D), 31 (Sham 28D). Mean ± SEM are shown as bars and lines. **(C)** Effect of Z944 (1µM, blue bars) on maximal rebound amplitudes recorded in current-clamp experiments in Sham WT and KO mice. Number of experiments were: 15 (Sham), 11 (Sham KO). Mean ± SEM are shown as bars and lines. **$p < 0.01$, ***$p < 0.001$, Wilcoxon test. **(D)** Maximal rebound amplitude recorded in current-clamp experiments after SNI in WT and KO mice. Mean ± SEM are shown as bars and lines. Dots show individual measurements. **(E)** Maximal rebound amplitude recorded in current-clamp experiments in males (black bars) and females (grey bars) naïve animals and after SNI in WT and KO mice. Mean ± SEM are shown as bars and lines. **(D, E)** Number of experiments were: 30 (Naïve), 61 (Sham), 36 (Sham KO), 46 (SNI 7D), 27 (SNI 7D KO), 48 (SNI 28D), 17 (SNI 28D KO). **(F)** Proportions of PKCγ neurons expressing cacna1h (CaV3.1) and cacna1g (CaV3.2), or cacna1i (CaV3.3) and cacna1g (CaV3.2) in the dorsal horn of lumbar spinal cords of Sham (n=4) and SNI (n=3) animals. *In situ* hybridizations were performed with the RNAscope technique (see Methods for details). **(G)** Effect of Z944 (1 µM) on maximal rebound amplitudes recorded in the current-clamp mode after SNI in WT and KO mice. Dots connected by a line are the data recorded from one neuron, before and during Z944 superfusion. Number of experiments were: 15 (Sham), 11 (Sham KO), 6 (SNI 7D), 4 (SNI 7D KO), 6 (SNI 28D), 4 (SNI 28D KO). *$p < 0.05$, **$p < 0.01$, ***$p < 0.001$, Wilcoxon test. **(G)**

### Supplementary Figure 3. Firing in PKCγ neurons.

**(A-B)** Cav3.2 ablation changed the resting resistance **(A)** and the peak amplitude **(B)** of the first action potential elicited upon depolarisation from -70mV in PKCγ neurons in Sham mice. Dots connected by a line are the data recorded from one neuron, before and during Z944 superfusion. Number of experiments were: 14 (Sham), 12 (Sham KO). *p < 0.05, **p < 0.01, Wilcoxon test. **(C)** Cav3.2 ablation did not modify the proportions of the firing patterns recorded in PKCγ neurons after SNI in WT and KO mice. **(D)** The mean intervals of the first action potentials during a 2s-long depolarisation in PKCγ neurons follow different relationships in WT and Sham KO mice. Mean ± SEM are shown as symbols and lines. The lower panel summarized the statistical differences between the intervals of the 1st, 3rd, 4th or 5th intervals with the interval of the 2nd action potential. #, §, £, & p<0.05 ; §§, && p<0.01 ; ####, ££££, &&&& p<0.001, Wilcoxon test. Numbers of experiments were: 37 (Sham 7D), 12 (Sham 7D KO), 40 (Sham 28D), 13 (Sham 28D KO).

### Supplementary Figure 4. Changes in IPSC of PKCγ neurons after SNI

**(A-C)** Inhibitory neurotransmission in PKCγ neurons did not change in Sham mice. **(A)** IPSC amplitudes and intervals were unchanged in PKCγ neurons in Sham mice, early (SHAM 7) and late (SHAM 28) after SNI. **(B, C)** IPSC amplitudes were similar in the dorso-ventral axis **(B)** and the medio-lateral axis **(C)** of PKCγ neurons in Sham mice. **(D,G)** The T-type channel blocker, Z944 (1 µM) did not change the IPSC amplitudes in PKCγ neurons of Sham WT **(D)** and KO mice **(E)**, early after surgery. The Z944 (1 µM) did not change the IPSC amplitudes in PKCγ neurons of Sham and SNI WT **(F)** and KO **(G)** mice, late after surgery. Graphs are cumulated distributions of the number of recordings indicated within brackets. Mean ± SEM are shown as symbols and lines.

### Supplementary Figure 5. Changes in EPSC of PKCγ neurons after SNI

**(A-C)** Excitatory neurotransmission did not change in PKCγ neurons of Sham mice. **(A)** EPSC amplitudes and intervals were unchanged in PKCγ neurons in Sham mice, early and late after SNI. **(B, C)** EPSC amplitudes were similar in the dorso-ventral axis **(B)** and the medio-lateral axis **(C)** in Sham mice. **(D)** Comparison of EPSC amplitudes in PKCγ neurons of SNI WT and KO mice. Graphs are cumulated distributions of the number of recordings indicated within brackets. Mean ± SEM are shown as symbols and lines. Statistical difference for ~20% of the events p<0.05 between appropriate distributions, Mann Whitney test.

**Supplementary Figure 1.**

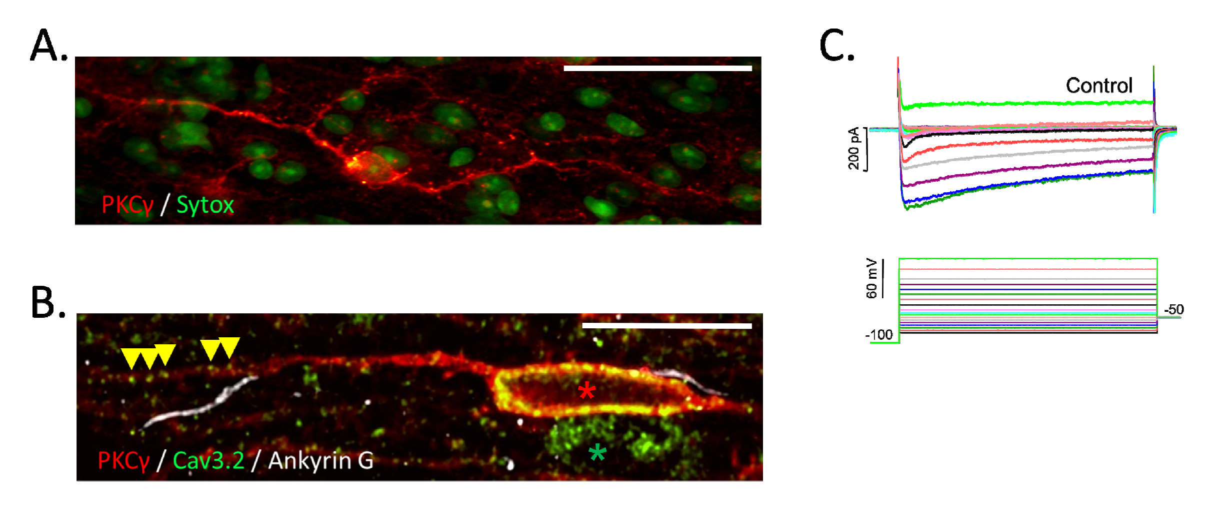

**Supplementary Figure 2.**

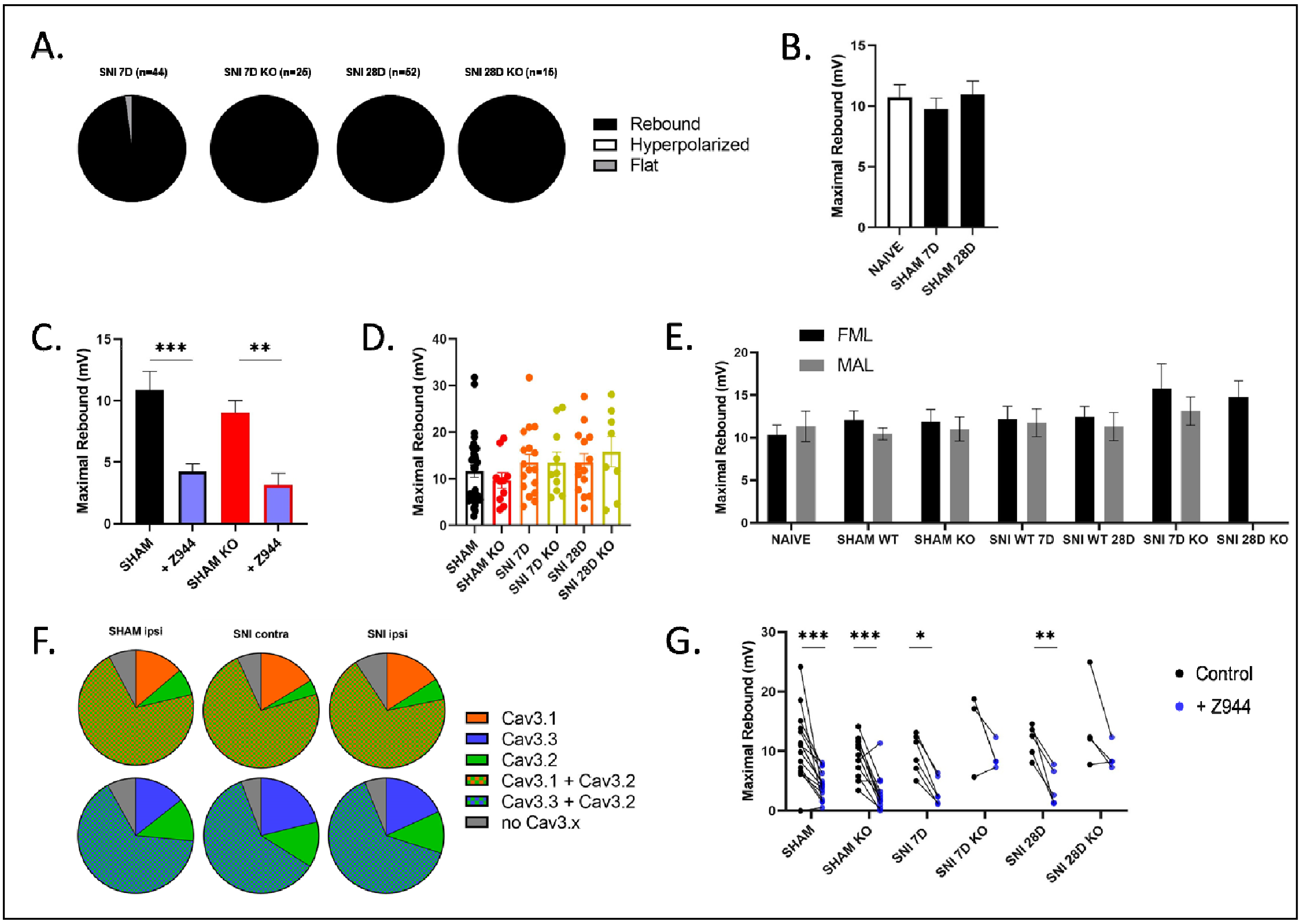

**Supplementary Figure 3.**

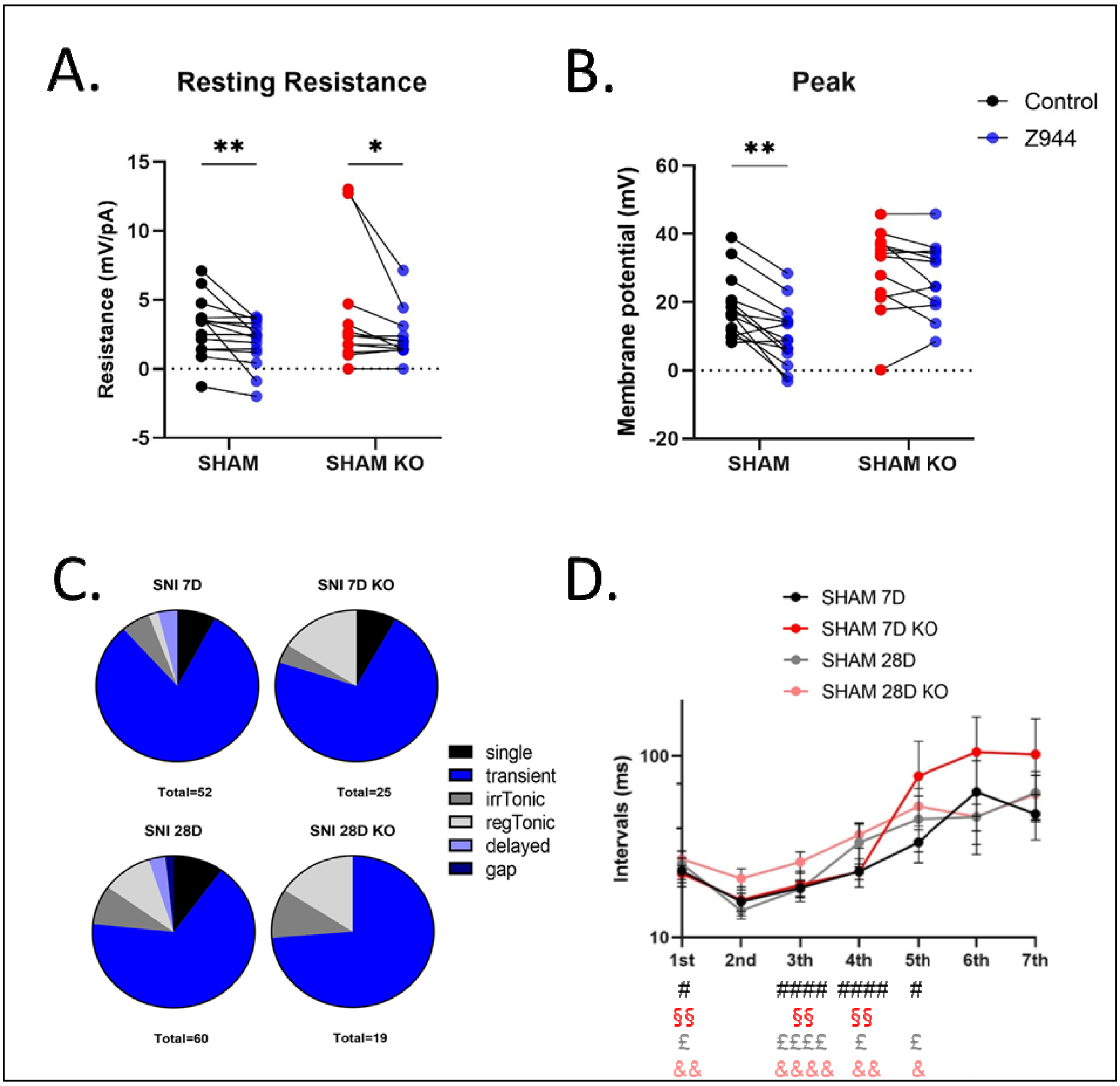

**Supplementary Figure 4.**

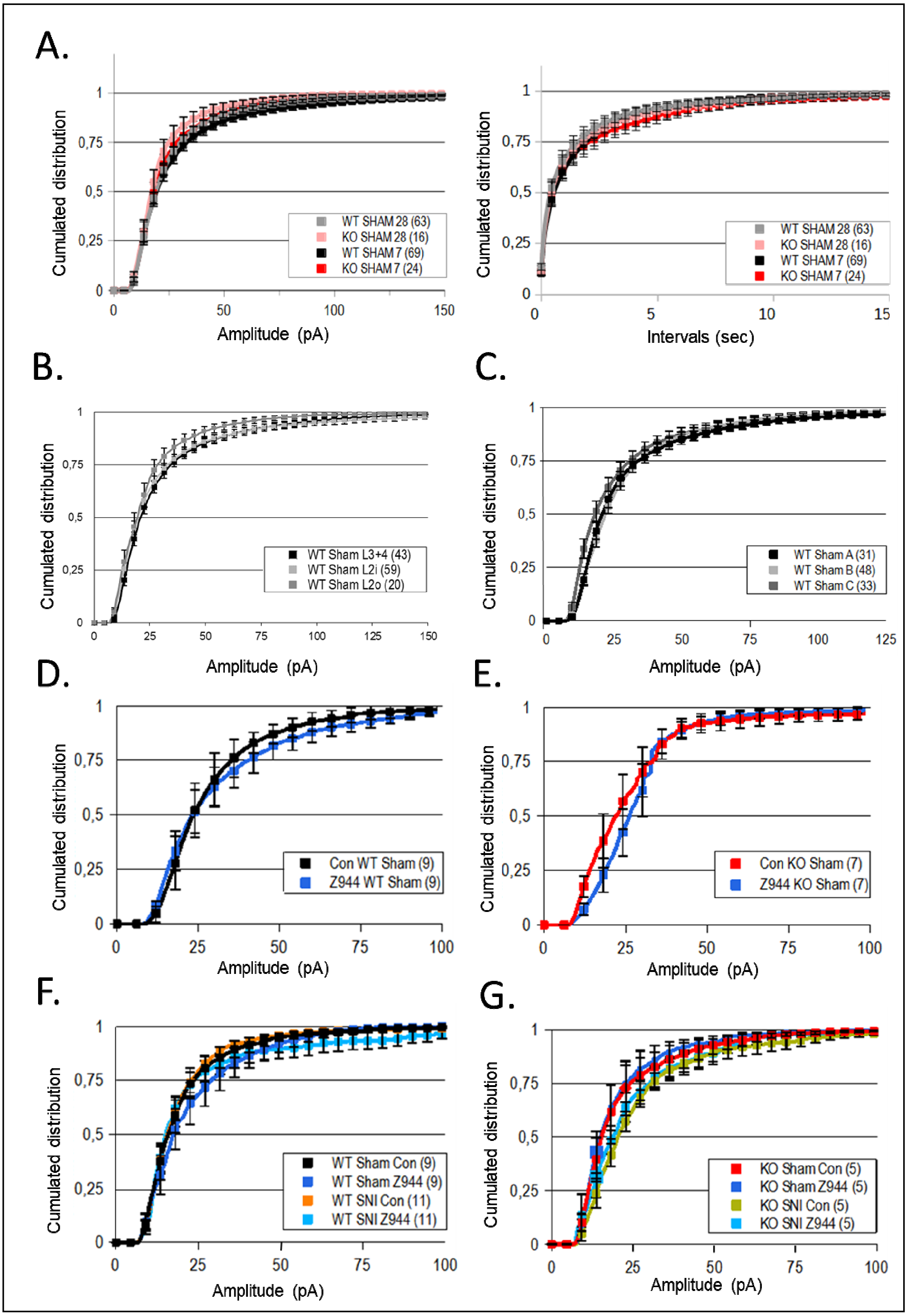

### Supplementary Figure 5.

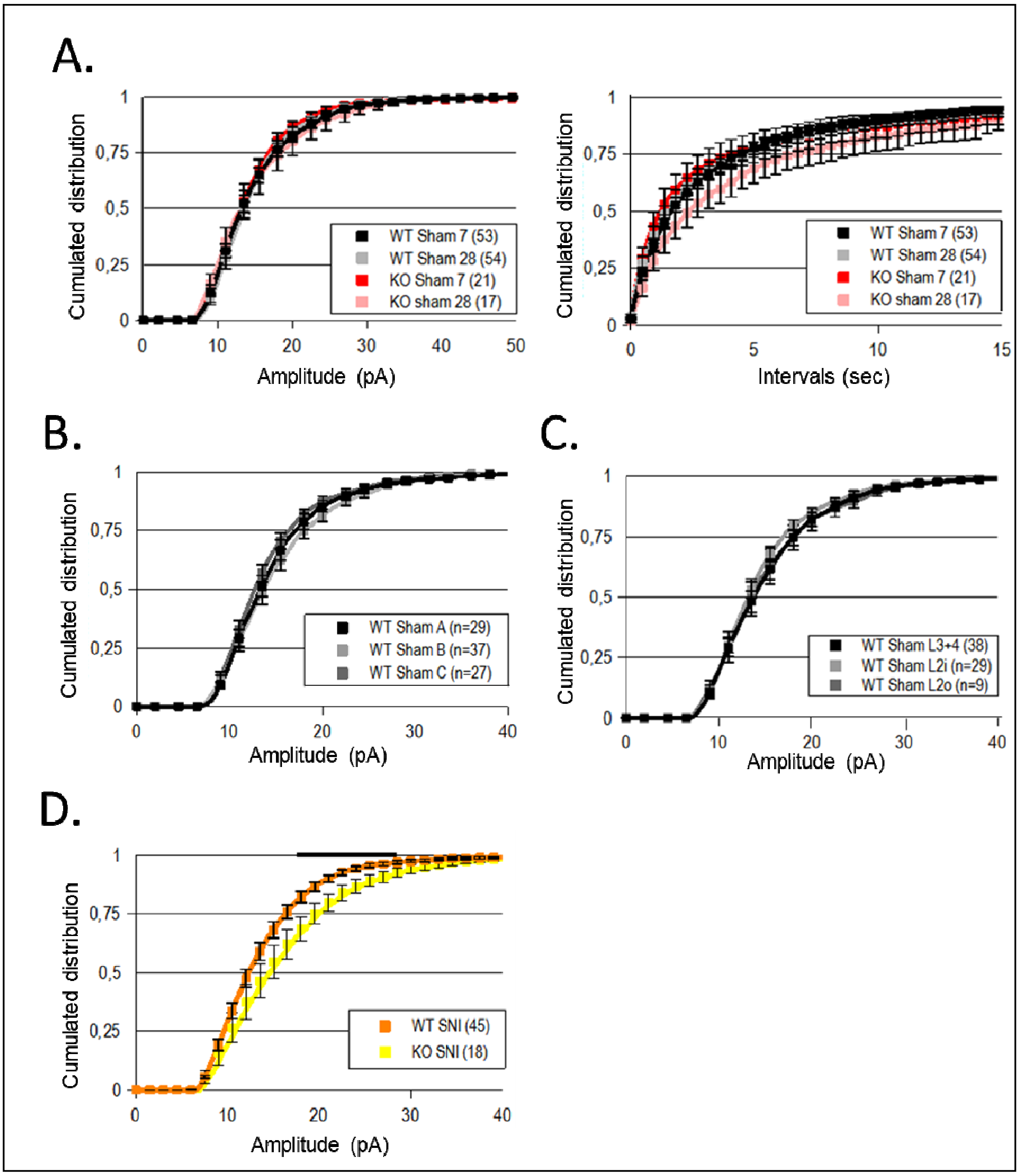
